# ORANGE: A Machine Learning Approach for Modeling Tissue-Specific Aging from Transcriptomic Data

**DOI:** 10.1101/2025.05.02.651895

**Authors:** Wasif Jalal, Mubasshira Musarrat, Md. Abul Hassan Samee, M. Sohel Rahman

## Abstract

Despite aging being a fundamental biological process which profoundly influences health and disease, the interplay between tissue-specific aging and mortality remains underexplored. This study applies machine learning on GTEx transcriptomic data to model tissue-specific biological ages across 12 different types of tissues and introduces an *age-gap* metric to quantify deviations from the chronological age. We use several modeling techniques optimized with three feature selection strategies: Pearson correlation, age-related differentially expressed genes, and tissue-enriched genes (expressed at least fourfold higher in a specific tissue). Among these, Pearson correlation combined with elastic net regression yields the best performance, with models achieving an average RMSE of 6.44 years and an R^2^ of 0.64. To quantify deviations from chronological age relative to the population, we train neural networks to regress predicted ages against chronological ages, and subtract their outputs from the predicted ages to calculate a metric which we call the *age-gap*. Age-gap statistics reveal significant tissue-specific aging patterns, identifying extreme agers and correlations between extreme aging and mortality. About 20% of subjects are found to exhibit extreme aging in one tissue, while 1% show multi-organ aging. Further analysis reveals that accelerated aging in specific tissues correlates with with greater risk of death from illness. These findings greatly emphasize the role of transcriptomics in aging research and its implications for health and longevity.

## 1 Introduction

Aging is closely tied to the onset of numerous diseases and the overall human health span. Recent advancements in molecular biology and machine learning have enabled researchers to delve deeper into the intricacies of biological aging, uncovering organ-specific patterns that diverge from chronological aging. A notable study by Oh and Rutledge et al. (2023) [1] utilized plasma proteomic data obtained via SomaScan [2] [3] assays to predict organ-specific biological ages and introduced the concept of *age-gaps*. These age-gaps, defined as the difference between an individual’s chronological age and the biological age predicted from proteomic data, correlate with health and disease states, offering a new lens on the interplay of aging and pathology. Inspired by this approach, we sought to explore the application of similar methodologies to transcriptomic data, specifically gene expression values measured as transcripts per million (TPM) from tissue samples in the GTEx dataset, incorporating subject sex as an additional variable. Building on the findings of Oh et al. [1] regarding the prediction of organ-specific age from the plasma proteome, we aimed to explore the potential of predicting tissue-specific age from the human transcriptome using machine learning models.

Despite the several epigenetic and proteomic studies in illuminating aging processes, the potential of transcriptomic data to predict organ-specific biological age remains underexplored. The GTEx dataset, a comprehensive resource of gene expression profiles across 54 human tissues, provides an opportunity to investigate this avenue. However, unlike plasma proteomics, transcriptomic data is inherently tissue-specific and requires a different approach to account for the unique expression patterns of each tissue. Additionally, challenges, such as limited sample sizes, variability in donor characteristics, and the lack of precise chronological age data in the public GTEx dataset complicate the modeling process. Addressing these challenges necessitates innovative preprocessing, feature selection, and predictive modeling techniques.

A recent study by Johnson and Krishnan (2023) [40] has associated the transcriptome with age and sex, utilizing transcriptomic data from RNA-seq samples to predict the age group of subjects within each sex group. However, it did not analyze the data on a tissue-specific basis, leaving unexplored the unique expression patterns across tissues. Furthermore, a multimodal approach to age estimation was recently applied to ovary and lung tissues from the adult GTEx dataset [41]. This approach combined transcriptomic data with additional modalities to improve prediction accuracy. For lung tissues, we improved upon the RMSE and R^2^ scores of their elastic net-based transcriptomic predictors. Remarkably, our models remain competitive even when compared to their multimodal ensembles making our simpler models more attractive and acceptable in this context. Previously, Ren and Kuan (2020) [27] proposed transcriptomic predictors for tissue-specific and cross-tissue ages using GTEx data. While their methodology provided a foundation for transcriptomics-based age prediction, our models are simpler, demonstrate improved performance in terms of RMSE and R_2_ scores on most tissues, and also generalize reasonably well to external datasets.

Furthermore, inspired by Oh et al. (2023) [1], we extend the scope of transcriptomic age prediction by incorporating the concept of *age-gaps* and related statistics along with their associations with mortality. The concept of the *age-gap*, which is the difference between an individual’s predicted biological age and their chronological age, has emerged as a significant metric in aging research. It offers insights into the rate at which an individual ages biologically, independent of the passage of time. A positive age-gap indicates accelerated biological aging, while a negative gap suggests a slower aging process. While Oh and Rutledge et al. (2023) [1] have presented the concept of a proteomic age-gap, most research has focused on the Brain Age-Gap (BAG) [14] [15] [16] which is associated with cognitive illness and dementia [17] [18] [19], as well as the Retinal Age-Gap (RAG) which is associated with mortality and reproductive aging [20] [21] [22]. Previously, Rutledge and Oh et al. (2022) [23] also applied the concept of age-gaps to omics data in general. In a recent study, Argentieri et al. (2024) [24] have also independently formulated the idea of a proteomic age-gap and have found it to be associated with mortality. Inspired by the existing body of research, we present the *transcriptomic age-gap* in an effort to enhance our understanding of the biological underpinnings of aging.

Building upon prior research, firstly we developed machine learning models to predict organ-specific biological ages using transcriptomic data from 12 selected tissues, focusing on tissues with sufficient samples and known correlations with mortality. Our methodology included robust feature selection methods, such as identifying age-correlated genes and analyzing differentially expressed genes with age, alongside advanced modeling techniques like bootstrap-aggregated Partial Least Squares (PLS) regression. By introducing a transcriptomics-based *age-gap* metric, we aimed to assess deviations in predicted biological ages and their associations with health outcomes and mortality. Analyzing our trained models, we found potential tissue-specific aging regulators. The results of our study highlight the viability of using transcriptomic data for organ-specific age prediction. Our models demonstrated competitive performance, achieving an average RMSE of approximately six years. We observed notable statistics in our analysis of age-gaps. Nearly 20% of the population in our study exhibited strongly accelerated aging in one organ, while a smaller subset, approximately 1%, was identified as multi-organ agers. These accelerated aging patterns were associated with mortality, conferring around 3 times the risk of death from intermediate or prolonged illness. Through this work, we aim to bridge the gap between transcriptomics and aging research, offering a new perspective on the dynamic interplay between biological age, health, and disease.

## 2 Methods

### 2.1 Dataset

We use the Adult Genotype-Tissue Expression (GTEx) v10 [5] dataset, which provides gene expression data measured in transcripts per million (TPM) across 54 tissue types from 948 adult subjects [10]. The dataset contains 19,788 tissue samples. This extensive dataset, collected from post-mortem tissue samples, includes gene expression data along with metadata [11] on age (binned into 10-year ranges), sex, circumstance of death characterized by the Hardy Scale (see Supplementary Section 1.1 for details on this scale) [47], and other clinical details, making it a valuable resource for understanding tissuespecific gene expression and genetic variations across a diverse human population.

### 2.2 Preprocessing and Initial Analysis

Initial analysis involved generating t-SNE plots of the tissue-specific gene expression values in TPM, resulting in distinct clustering of samples corresponding to individual tissues (Figure S5).

#### 2.2.1 Tissue Selection for Predictive Modeling

Since the Adult GTEx does not contain samples of every tissue type from each test subject, and the aging of all organs does not have a strong correlation with mortality, we had to limit our study to a set of tissues that had a significant number of samples and are known to be correlated with mortality [8] [9]. The selected tissue types were liver, aorta, coronary artery, brain cortex, brain cerebellum, heart atrial appendage, subcutaneous adipose, lung, sun-exposed skin, tibial nerve, sigmoid colon, and pancreas. We plotted t-SNE plots for the samples for individual tissue types and observed no distinct clustering based on age range. However, in certain cases, clustering was evident when categorized by the Hardy Scale ratings (Figure S6).

#### 2.2.2 Optimal fixed-point interpolation for age-ranges

To compensate for the absence of precise ages of the subjects in the dataset, we opted to find optimal points within each age range using an exhaustive search approach, rather than just taking the midpoint of each range. Each age range was divided into three points using the formula:

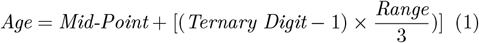

Iterating through 3^6^ permutations on six age ranges for each tissue type, based on the RMSE and *R*^2^ scores achieved by each permutation, we found the most commonly-occurring optimal permutation of ternary digits to be 222100. As expected from a Gaussian distribution (Figure S1), the optimal point for each age range deviated from the range mid-values towards the population mean (see Supplementary Section 10).

### 2.3 Predicting Age from Gene Expression

To develop tissue-specific aging models, we grouped the gene expression data by tissue type, and then performed all training and testing on tissue-specific data only. We initially partitioned each tissue’s dataset into training and testing subsets, ensuring that all samples with a Hardy Scale rating of 1 (see Supplementary Section 1.1) were allocated to the testing sets, since the age at death for subjects who died of unnatural causes such as accident or suicide may not be reflective of their tissue health. We assigned the remaining subjects to the training and testing subsets randomly to get an overall train-test ratio of approximately 80:20.

#### 2.3.1 Feature selection by identifying age-correlated tissue-specific genes

We selected features for tissue aging models applying three different methods. In the first method, we conducted a correlation analysis on the columns of expression values for each tissue, and selected only the columns exhibiting a Pearson correlation greater than 0.2, thereby retaining features that demonstrate a significant linear relationship. In the second method, we chose genes that were differentially-expressed with age, using the package PyDESeq2 [38], which is a Python implementation of the Bioconductor package DESeq2 [4]. We fixed the optimal log_2_(*fold-change*) threshold for each tissue using an exhaustive search approach, and selected the genes that showed the highest fold-changes across age groups. In the third method, we followed an approach almost identical to the one used by Oh et al. [1] to identify organ-enriched plasma proteins. A gene is organ-enriched if it is expressed at least four times higher in a single organ compared to any other organ [42]. We define tissue-enriched genes as genes that are expressed at least four times higher in a single tissue than in any other tissue. The log_2_(*fold-change*) of normalized read count of each gene in each tissue was determined using the PyDESeq2 [38] package, and each gene was assigned as a feature to the tissue in which its fold-change was the greatest.

#### 2.3.2 Feature transformation and learning algorithms

In order to stabilize variance and make the gene expression data more Gaussian-like, the Yeo-Johnson power transformation [7] was applied to the expression value columns.

Utilizing the selected gene expression columns for each organ, along with sex (encoded as 0/1) and the optimal fixed-point within each age range from the age column as target variables, we trained linear predictors. The results were averaged through bootstrap aggregation to enhance model robustness and predictive accuracy.

For modeling tissue-specific age, we employed 20× bootstrapped predictors of several types. Following the method of Oh et al. [1], we tested LASSO regression, which prevents overfitting using L1 regularization, and elastic net regression, which combines L1 and L2 regularization. We also applied Partial Least Squares (PLS) regression [39], which projects the features and targets onto a shared low-dimensional space, an approach well-suited for high-dimensional and collinear predictors. In parallel, we experimented with Support Vector Regression (SVR) [61] using linear kernels, which approximates the target function within a tolerance of *ϵ* while maximizing the margin, and with random forest regressors [62], which aggregate predictions from multiple decision trees.

The mathematical formulations of the learning methods are provided in Supplementary Sections 2 and 3.

#### 2.3.3 Age-gap estimation

Following the prediction of tissue-specific ages, we calculated a regression for the predicted ages of tissue samples against the chronological ages of their respective subjects using a neural network containing two fully-connected layers (1 × 32 and 32 × 1) with batch normalization and ReLU activation layers in between, thus yielding a *ŷ* value for each test sample. We used this approach instead of a simple linear regression to account for any possible nonlinearity in the underlying relationship between gene expression and age. Inspired by the concepts presented by Oh et al. [1], we defined a metric called the *age-gap*, by subtracting the *ŷ* value from the predicted age. This age-gap serves as an indicator of a tissue sam-, ple’s aging relative to its counterparts within the same age group.

### 2.4 Result Interpretation and Downstream Analyses

#### 2.4.1 Evaluation metrics

We evaluated the predictive performance of our models (i.e., predicted age vs. chronological age) using two metrics: the RMSE (Root mean-squared error) and the *R*^2^ (coefficient of determination). To choose the optimal model for downstream analyses, we also considered training speed as a metric.

#### 2.4.2 Leave-*P*-out training-testing for better downstream analyses on age-gaps

We aimed to generate a larger set of predicted age-gaps across all subjects, in order to examine the correlation between extreme aging and circumstance of death (characterized by Hardy scale ratings) through a conditional probability analysis. To compensate for the small subject count of the dataset, apart from the train-test splitting strategy presented in Section 2.3, we also adopted another strategy, where after allocating the samples with Hardy Scale ratings of 1 directly to the test set, we selected *P* of the shuffled samples from each tissue to perform a leave-*P*-out train-test splitting in a sliding window mechanism, where *P* equals 5% of the total number of samples from that tissue. This strategy ensured that we covered each sample in the tissue’s dataset. Each 5% window was used as the testing set, on which predictions were performed by training on the rest of the samples. This enabled us to perform a prediction on every single sample and ultimately have a larger set of predicted age-gaps, on which we could perform better downstream analyses. The samples with Hardy Scale ratings of 1, that we had initially separated for testing only, were appended to the last testing window of each tissue’s dataset. For this leave-*P*-out workflow and all downstream analyses, we used a training architecture that involved feature selection based on absolute Pearson correlation, followed by bootstrap-aggregated PLS regression without cross-validation. This approach provided the best balance between prediction accuracy and execution speed (see Supplementary Tables 9 and 10). We ran 20 cycles of this leave-*P*-out train-test exercise by varying a random seed each time that affected only the initial shuffling that we performed to ensure class homogeneity.

#### 2.4.3 Conditional probability analysis

For each tissue type, we made dot plots of the age-gaps calculated from our predictions through the leave-*P*-out exercise and tried to fit them under normal distributions. At the two extremes of such distributions, we identified groups that we refer to as ***extreme negative agers*** and ***extreme positive agers***, defined as those falling below 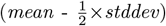 and above 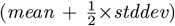,. respectively.

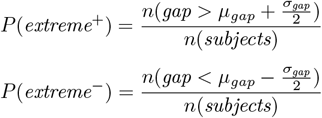

Conversely, individuals situated near the center of the distribution, approximately within 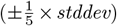 around the mean, were categorized as ***average agers***.

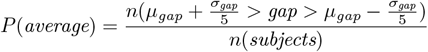

To explore the contributions of extreme age-gaps in disease and mortality, we considered the conditional probability ratios –

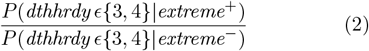

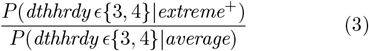

On the other hand, to determine how age-gaps may relate to subjects who died unnatural deaths but may have been otherwise healthy (*dthhrdy* = 1), we considered the following ratios –

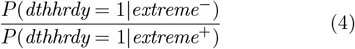

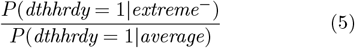

In the above equations, *dthhrdy* is the Hardy Scale rating (see Supplementary Section 1.1) of the circumstance of subjects’ deaths.

#### 2.4.4 Code Environment and Availability

All the predictors were trained and tested using the Python package scikit-learn [6]. The hyperparameters *α* (L1 regularization parameter for LASSO and elastic net), *C* (regularization parameter for SVR), and *n*_*estimators*_ (number of trees in random forest regression) were all tuned with four-fold cross-validation using the GridSearchCV tool from scikit-learn. Fully-connected layers used in age-gap estimation were implemented using the PyTorch [43] library. We have made the code available through a GitHub repository (https://github.com/wjalal/ORANGE/).

Further details regarding our experimental setup are provided in Supplementary Section 4.

## 3 Results

### 3.1 Model Performance

We observed varying model performance per tissue in terms of prediction accuracy and training speed in the various combinations of the methods we used for feature selection and training (Supplementary Tables 9 and 10). In general, the selection of characteristics by absolute Pearson correlation performed better than the selection by tissue-specific DEGs (differentially-expressed genes). Elastic net predictors had the best RMSE and *R*^2^ scores on average, with LASSO and PLS regressions closely following it. PLS regression without cross-validation performed the fastest while also predicting sample ages with competitive accuracy. The PLS approach greatly reduces the dimensionality of the dataset and performs much faster than the Ordinary Least Squares (OLS) regressors Lasso, Ridge, and Elastic Net which employ L1 and L2 regularization. PLS regression offered the best balance between training speed and accuracy. It also achieved competitive performance without the use of cross-validation in training, without which the OLS regressors could not perform that well (Supplementary Table 1).

When tested on common datasets, our PLS regression models also performed well compared to models developed in other studies (Supplementary Table 11). The RNAAgeCalc models developed by Ren et al. (2020) [27], which were trained on the GTEx v6 data, did not generalize well when we used them to predict the age of GTEx v10 samples that were not present in GTEx v6. However, our models, when trained on the portion of GTEx v10 data which overlaps with GTEx v6, achieved better average RMSE and *R*^2^ scores of 6.99 years and 0.49 respectively on test samples. Note that, the model was tested on GTEx v10 samples that were not present in GTEx v6. Our elastic net and PLS regression models for lung tissue aging also performed noticeably better than the gene expression-based aging models used by Ribeiro et al. (2024) [41] in their study on multimodal modeling of human aging. In fact, despite having a more constrained setup and having gene expression data as the sole modality, on a common testing set, our transcriptomic aging models exceeded the performance of all of their models except the epigenomic ones (methylation-based) and performed significantly better than their gene expression-based models (both elastic net and gradient boosted trees).

To explore the generalizability of our models, we tested our GTEx-trained elastic net models on independent datasets (see Supplementary Section 6 for methodological details). Our lung aging model achieved an RMSE of 7.79 years and an *R*^2^ of 0.36 on a set of 46 normal tissue samples from the Cancer Genome Atlas Lung Adenocarcinoma (TCGA-LUAD) [12] dataset. Combining the results produced by our subcutaneous adipose, liver, lung, and kidney cortex models, we achieved RMSE and *R*^2^ scores of 8.16 years and 0.32 respectively on a total of 30 samples from transcript-level RNA-seq data from the Human Protein Atlas project [13].

### 3.2 Gene TPMs can model organ age

Our bootstrap-aggregated linear models could capture clear linear relationships between the TPM values of differentially-expressed genes measured from tissue samples and the subjects’ chronological ages (Figure 3). Our best models achieved an average (weighted by number of samples analyzed per tissue type) RMSE of 6.44, and *R*^2^ of 0.64 for predicted ages.

#### 3.2.1 Distribution of extreme agers across the population

We observed that most individuals had varying age-gaps across organs (Figure 2a). Analyzing the statistics of the predicted age-gaps among 531 subjects led us to some interesting findings. We observed low average correlation between the age-gaps of different tissues of a subject (Figure 2b), with an average pair-wise Pearson correlation of 0.21 across all tissue types. However, no negative correlation was observed, and in general, most of an individual’s age-gaps (across tissue types) centered around an average (Figure 2a), due to a low correlation. About 26% of all subjects were found to have extreme aging (absolute value higher than two standard deviations) (Figure 2c).

**Figure 1.**
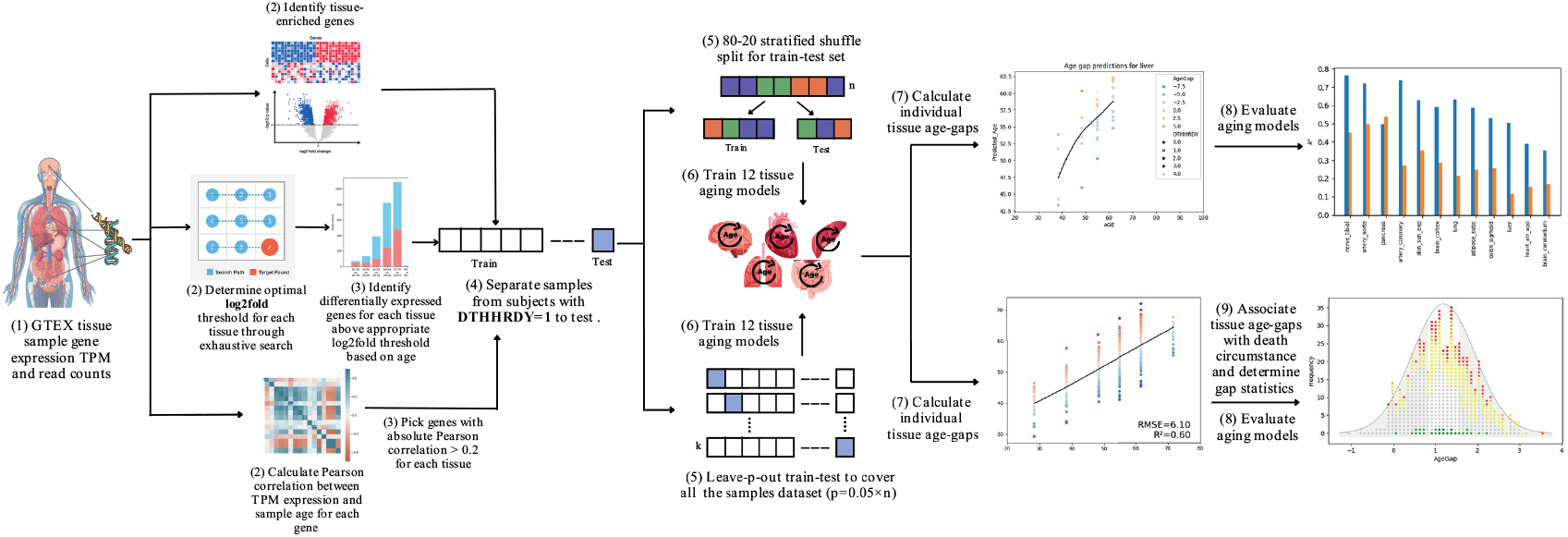
Study Methodology. The study utilized GTEx v10 data to predict tissue-specific biological aging from gene expression profiles. Key tissues were selected based on sample size and mortality correlations, and machine learning models were trained on age-correlated features for each tissue. An *age-gap* metric was introduced to quantify deviations in predicted tissue age from chronological age, revealing extreme agers in specific tissues. The analysis highlighted patterns of accelerated aging in a small subset of subjects, providing insights into tissue-specific aging dynamics and their connection with mortality.

**Figure 2.**
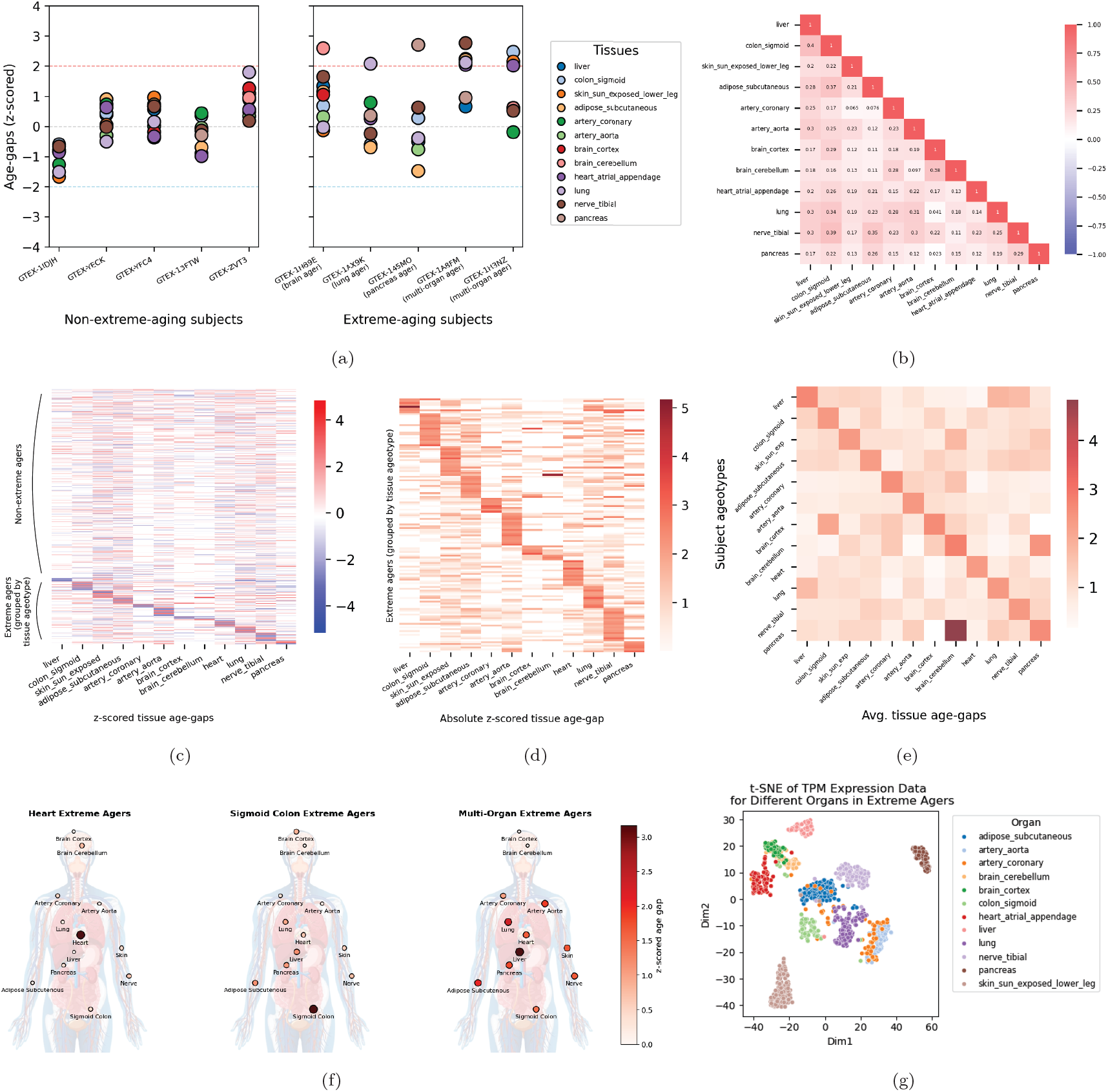
Distribution of extreme agers across the population. **(a):** Extreme agers (at least one tissue’s z-scored age-gap *>* 2) and non-extreme agers. Five randomly-sampled subjects without extreme aging are shown; all appear to have varied tissue age-gaps that are slightly correlated and thus scattered around a mean. About 26% of all subjects were found to have extreme aging, and most of them (21% of all subjects) exhibited extreme aging in only one organ (three example subjects shown), while only 0.94% of the subjects showed multi-organ aging (two samples shown). **(b):** Heatmap of pairwise correlation between the age-gaps of each tissue type, 0.21 on average. **(c):** Heatmap of z-scored age-gaps of all subjects, partitioned into non-extreme agers and extreme agers grouped by tissue ageotype. **(d):** Heatmap of absolute z-scored age-gaps of extreme agers only, grouped by tissue ageotype (tissue with highest age-gap). Most agers show relatively accelerated aging in only one organ, by a notable margin. **(e):** Average age-gaps in each tissue (column-wise) of the subjects of each tissue ageotype (row-wise). Almost each ageotype has the highest average age-gap in the corresponding tissue by a notable margin. Note that the subjects with extreme-aging brain cortex tissues also have a higher average cerebellum age-gap, indicating that there may be a common group of brain agers among them. **(f):** Average tissue age-gaps of subjects with extreme aging in heart atrial appendage and sigmoid colon tissues shown as examples. Multi-organ agers show extreme aging across tissues, whereas most others show extreme aging in only one tissue along with low absolute age-gaps in other tissues. **(g):** t-SNE plot of tissue gene expression values (TPM) of extreme agers. Expression values of the same tissue cluster together, as do those of related tissues.

**Figure 3.**
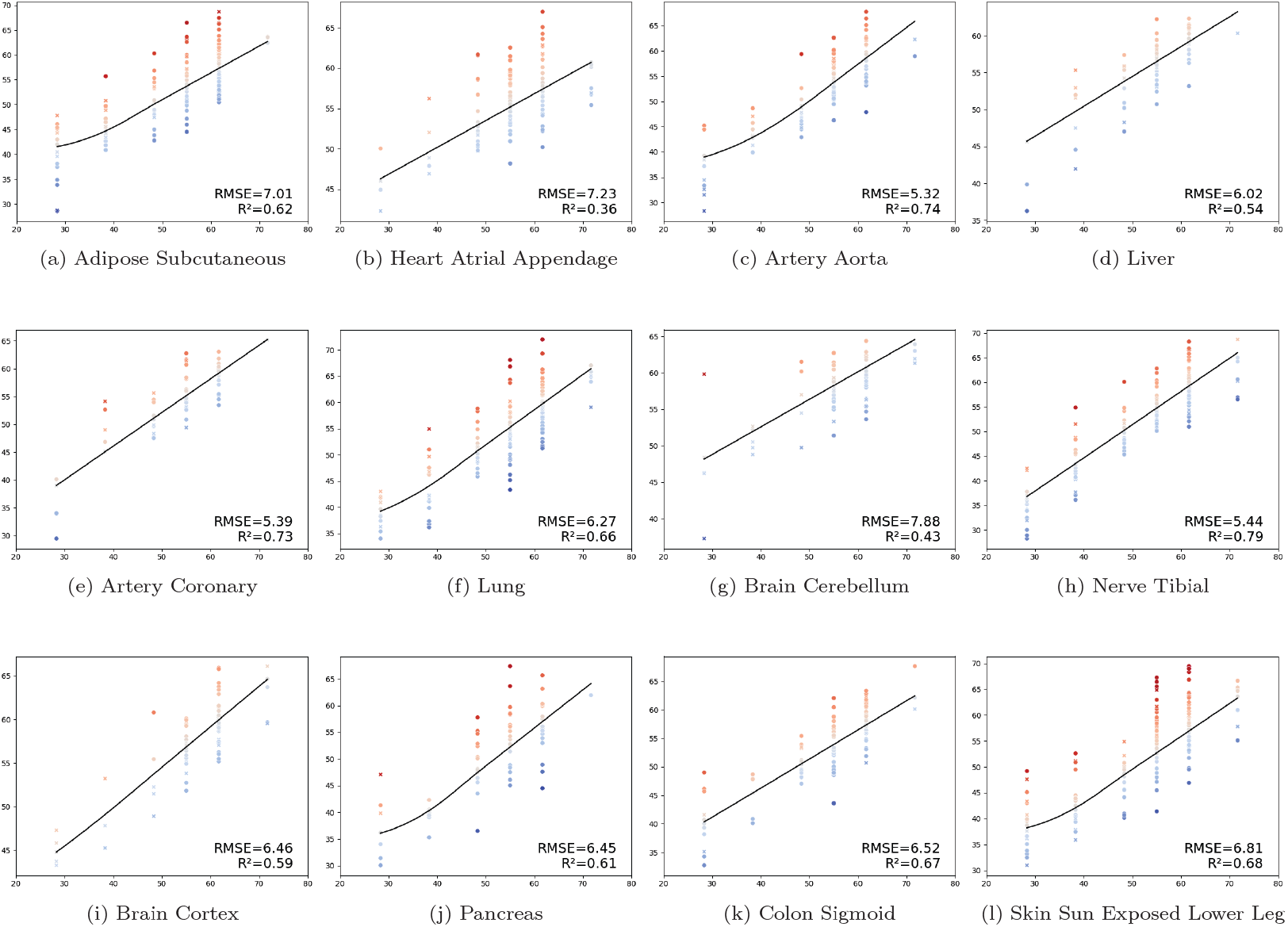
Modeling tissue age with gene TPM. Model performance across various tissue types, measured in *R*^2^ and RMSE, using Pearson correlation-based feature selection and elastic net predictors. Each plot shows the testing set samples of the corresponding tissue as dots whose positions along the plot’s x-axis and y-axis represent the corresponding sample’s chronological age and predicted age respectively. The line of fit in each plot is estimated by neural networks described in Section 2.3.3, while the vertical distance between each dot and the line of fit represents the corresponding sample’s age-gap.

These gene expression values of the samples from these subjects clustered together by tissue, and the clusters of the individual tissues mostly remained separate from each other (Figure 2g). Interestingly, 21% of the subjects exhibited extreme aging in only one organ (Figure 2d), while only 0.94% of the subjects showed multi-organ aging (extreme aging in 3 or more tissues), suggesting that most cases of extreme aging are organ-specific (Figure 2e, 2f). With our novel approach, which is different from existing ones in the literature, we effectively reached conclusions that are in line with the current knowledge base [1]. In addition, we were able to identify some new relevant insights which call for further biological validation, thereby opening new avenues for research.

### 3.3 Age-gaps are associated with circumstance of death

Dot plots of each subject’s tissue-specific age-gap or maximum observed age-gap across the selected 12 tissue types appear to fit normal distributions (Figure 4(a)-(l)). We could observe differences in the distribution of the five Hardy Scale death circumstances within the various regions under the normal curves (*extreme positive agers, extreme negative agers*, and *average agers*).

**Figure 4.**
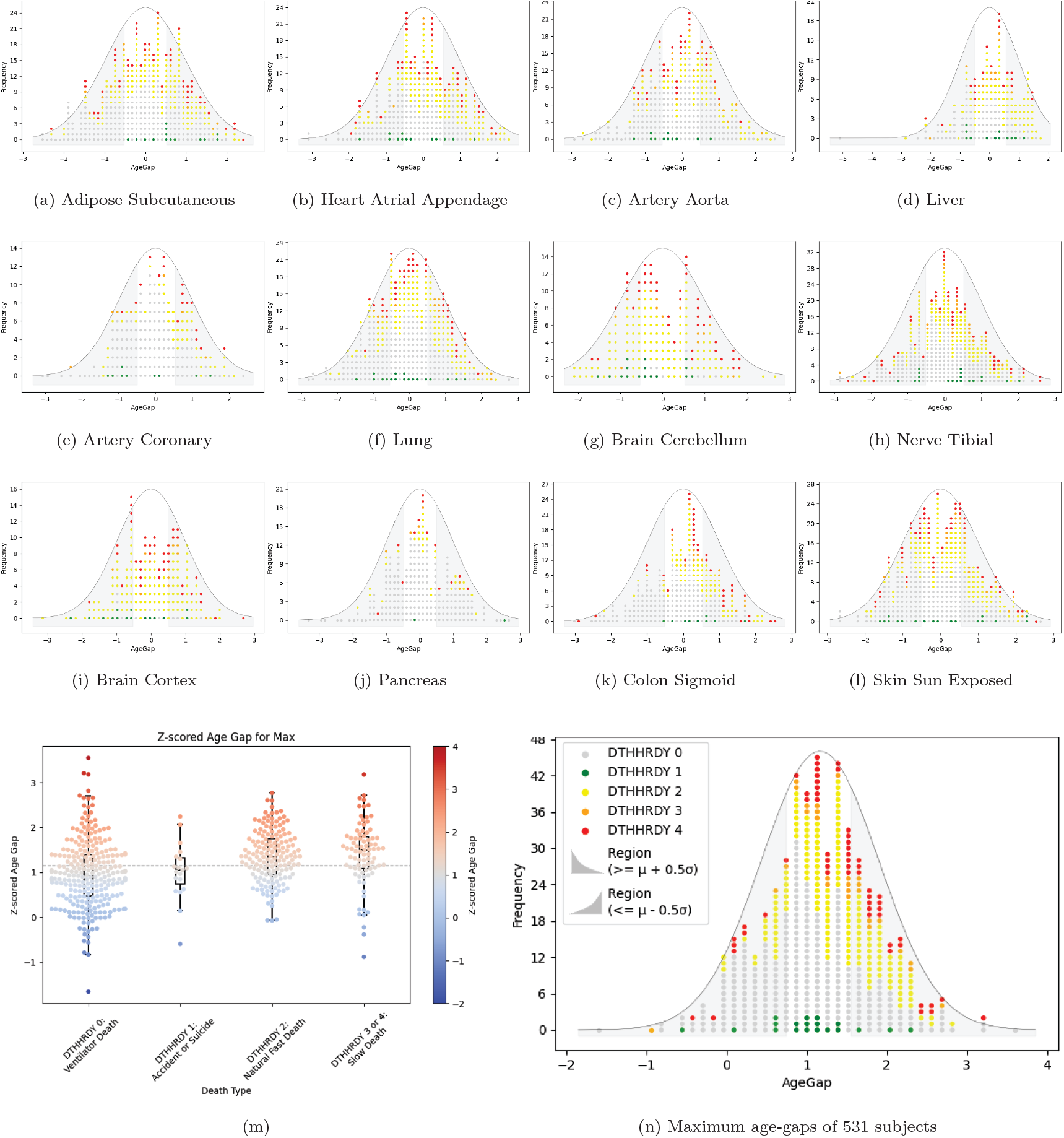
Tissue age-gaps of vital organs correlate to cause of death. **(a)-(l):** Distribution of age-gaps calculated by predicting ages of the samples of a tissue type. The age-gaps fit normal distributions. Regions of interest in the tails of the curve represent the conditions for ***extreme negative agers*** and ***extreme positive agers***, defined as those falling below 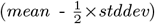 and above 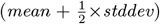, respectively. **(m):** Distribution of maximum age-gaps of 531 subjects with samples of more than 5 tissue types, compared across *dthhrdy* classes. Subjects who died of prolonged illness (*dthhrdy* 3 or 4) had their maximum (worst-case tissue) z-scored age-gaps distributed above the mean. **(n):** Distribution of maximum age-gaps of 531 subjects with samples of more than 5 tissue types. The legend in *(n)* applies to all the plots *(a)-(l)* and *(n)*, to indicate the conditional probability of finding each *dthhrdy* type among the two types of extreme agers.

#### 3.3.1 Tissue age-gaps of vital organs correlate to cause of death

Across all 12 tissues that were analyzed, we found that *extreme positive agers* were significantly more likely to have died with *dthhrdy* = 3 or 4 (intermediate death with a short terminal phase, or slow death after a long illness; both classified as ill subjects), than *extreme negative agers*, and moderately more likely compared to *average agers* (Figure 4m). We argue that the maximum age-gap that a subject has across the selected 12 tissue types, could be an indicator of their worst-aged organ, and possibly be associated with the subject’s cause of death. From the conditional probability analysis of *dthhrdy* (Hardy Scale rating for circumstance of death) distribution by age-gap (Figure 4n, Supplementary Table 8), we could conclude that, *extreme positive agers* were 2.96 times as likely to have died with *dthhrdy* = 3 or 4 (intermediate death with a short terminal phase, or slow death after a long illness; both classified as ill subjects), than *extreme negative agers*, and 1.24 times as likely as *average agers* (based on Eqns 2 and 3). It was also observed that *extreme negative agers* were 1.27 times as likely to have died with *dthhrdy* = 1 (death due to accident, blunt force trauma, or suicide) as *extreme positive agers*, and 0.59 times as likely as *average agers* (based on Eqns 4 and 5), suggesting that subjects whose tissues had decelerated aging relative to their chronological age were more likely to have died due to unnatural causes rather than diseases.

### 3.4 Identifying important regulators for tissue aging

The highest-magnitude coefficients in our linear models for tissue age prediction offered insights into several genes that are known to specifically regulate aging in the corresponding tissues (Figure 5) as well as across multiple tissues. A total of 17 genes were identified as important age regulators across multiple tissues, while a total of 583 were observed as regulators of tissue-specific age.

**Figure 5.**
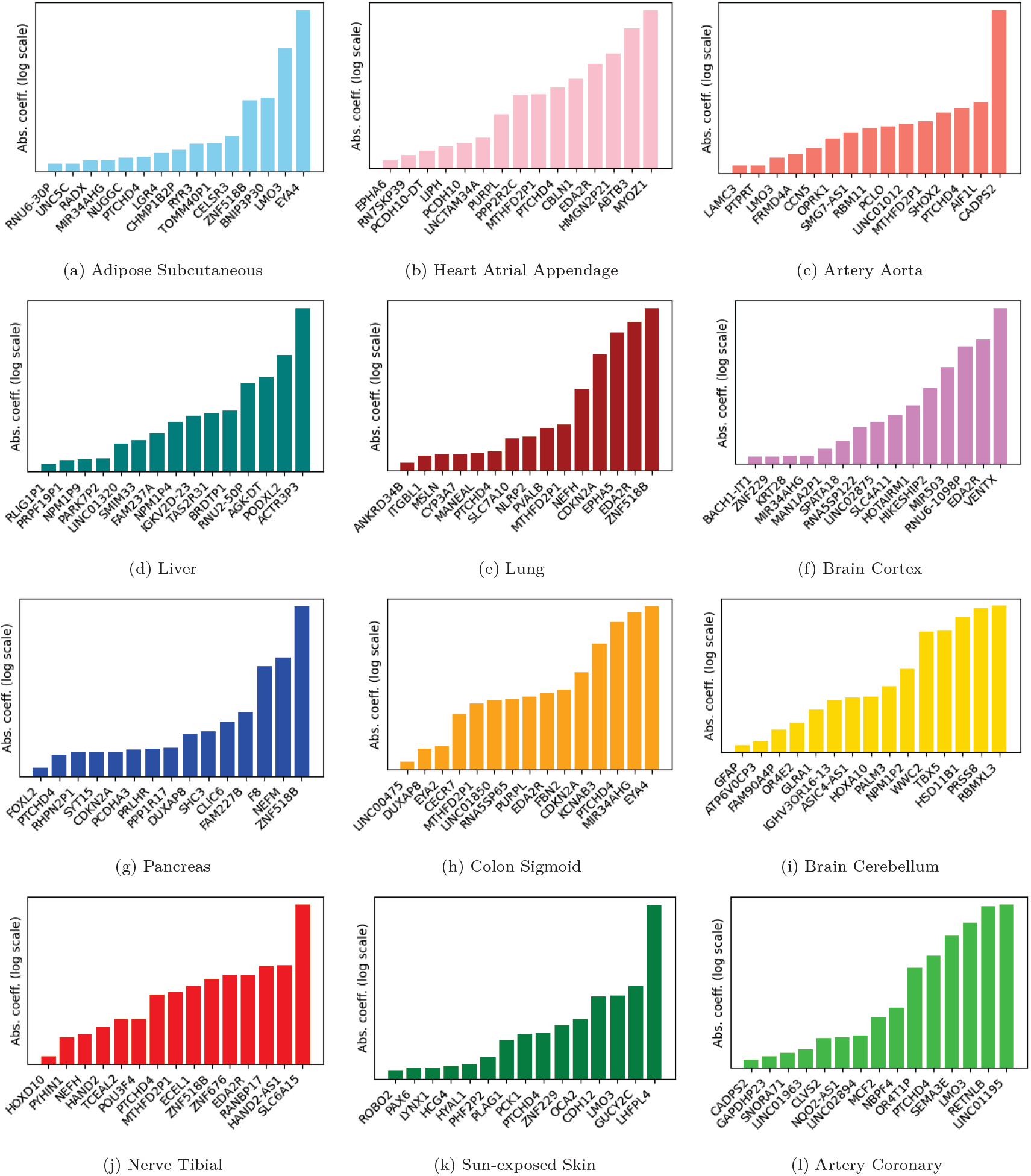
Top 15 Absolute Coefficients for Various Tissues. Plots show the significant features contributing to model predictions for each tissue.

#### 3.4.1 Tissue-independent or multi-tissue determinants of aging

Our aging models for each of the analyzed tissues, except for liver and brain, assigned high positive weights to the expression of PTCHD4, a gene which interacts with pathways involved in DNA repair and cell cycle regulation, particularly in the context of the p53 pathway [44] [45]. PTCHD4-AS, a related long non-coding RNA, has shown potential in enhancing DNA repair and possibly activating checkpoints that could protect cells from accumulating damage-a factor that could influence aging by either supporting cell survival or promoting senescence to prevent malignancy [25].

EDA2R, which has been described as the most remarkable and ubiquitous aging-related transcriptional hallmark [29] and a general marker of aging across tissues [27], was weighted significantly in our brain cortex, heart, adipose, lung, pancreas, and colon aging models. The expression of EDA2R in adipose, heart and lung was also independently found to be connected with aging [28] and cellular apoptosis, suggesting that it is an inducer of apoptosis [26]. LMO3, whose expression has been found to increase with age in adipose tissues, bears major weight in our artery, heart, adipose and skin tissue aging models. ZNF518B, which bears negative weight in several of our models, has been observed to be consistently downregulated with age in both humans and mice [33]. Interestingly, the pseudogene MTHFD2P1 is positively correlated with age in several of our tissue aging models, which prompts further biological studies to explore its role in aging, particularly because the related protein-coding gene MTHFD2 is called a potential oncogene [34] and its corresponding enzyme has been shown as an aging-associated factor in cancer [35].

MIR34AHG, the host gene for MicroRNA34-a, which is an important mediator of inflammaging [32], was also identified as a major contributor to age in several of our tissue aging models. Studies have shown that the expression of MicroRNA-34a increases with age [30] and it regulates cardiac aging and function [31], which has been reflected in the positive coefficients assigned to its expression in our models for heart atrial appendage and other tissues.

#### 3.4.2 Tissue-specific determinants of aging

One of the genes that our subcutaneous adipose tissue aging model identified as a key determinant of aging in the tissue was ZMAT3, whose upregulation has been associated with senescence in subcutaneous adipose [36], activation of the p53/p21 pathway, and the inhibition of adipogenesis. Another gene called OCA2 bears a negative weight in our skin aging model, thereby aligning with its established role in melanin production and skin pigmentation, reinforcing previous findings linking melanin levels to skin aging and cancer risk [59] [60].

The RNA gene, PURPL, which promotes tumorigenicity in colorectal cancer [37], was assigned a high coefficient by our sigmoid colon aging model. In addition, the model highlighted DUXAP8, a pseudogene that contributes to colorectal cancer progression by inducing epithelial-mesenchymal transition [48].

In the brain, our cerebellar aging model identified GFAP as a significant gene associated with aging. This gene encodes the glial fibrillary acidic protein, which is an intermediate filament protein found in astrocytes. Elevated GFAP levels in cerebrospinal fluid have been proposed as biomarkers for neurodegenerative diseases, such as Alzheimer’s disease [50], while GFAP has also been found to be elevated in the cerebral cortices of autistic subjects [49], suggesting that the encoding gene is a marker for neurodegeneration and brain-aging. Additionally, our brain cortex aging model identified MIR34AHG as a key factor. MIR34A, a protein encoded by MIR34AHG, has been shown to modulate inflammatory molecules involved in post-stroke recovery and affect blood-brain barrier permeability [52]. Its potential role in Alzheimer’s disease, suggested by a study that finds it overexpressed in the cortex [51], further supports its relevance in brain aging.

Our lung aging models identified MSLN, the protein-coding gene for Mesothelin, as a regulator of aging, and its relevance is supported by several studies that link the expression of Mesothelin with human lung cancer [53] [54] [55]. EPHA5, another gene whose expression has been associated with the development of human lung adenocarcinoma, was also found to be an important regulator of lung aging by our model. On the other hand, NLRP2, which has been identified as an antioncogene which inhibits the proliferation of lung adenocarcinoma cells [56], was assigned a negative weight by our model.

In the pancreas, an important determinant of aging identified by our model is the tumor suppressor gene CDKN2A, which is a well-known catalyst in the development of pancreatic ductal adenocarcinoma [57]. Our pancreas aging model also assigns a strong weight to PRLHR which is the encoding gene for the prolactin-releasing hormone receptor. Its role in pancreas aging is supported by literature that establishes prolactin as a promoter of pancreatic cancer progression [58].

## 4 Discussions

In this study, we have developed a mechanism to model tissue-specific biological aging from the transcriptome of post-mortem tissue samples. Our models can estimate the biological age of tissue samples from the expression of identified genes measured as TPM (transcripts per million) through RNA-seq, model tissue-specific aging profiles, and identify transcriptomic markers of aging.

Although the GTEx is widely regarded as the largest dataset for tissue-specific human transcriptomic data, there were many limitations in our study owing to the restrictions of the dataset. Despite having a large number of tissue samples, the number of individual subjects is around 1000, which is relatively small for analytics at the individual level. The open-access Adult GTEx only has limited data on donor phenotypes, which are sex, 10-year age bracket, and Hardy scale rating. Unbinned age, race, weight, smoking status, diabetes status, and other disease-related de-identified donor phenotypes are available in the protected-access dataset, which potentially holds future directions for our work. Despite using a 10-year age bracket, our age prediction models had RMSE scores around 6, which is comparable to or even better than models that used exact ages [27] [41]. It is also worth mentioning that the GTEx project collects samples only from tissue sites that it classifies as *non-diseased*; so the inclusion of diseased tissues could change the picture of modeling tissue-specific aging and disease risk. The project does however include samples from subjects with Hardy scale ratings defined as death after prolonged disease. Future studies on this topic should focus on tissue-specific disease prediction and modeling.

We had to restrict our study to a specific set of tissues, because not all the tissue-types in the dataset have sufficient samples, and most tissues are not strongly correlated with mortality. We developed prediction models for reproductive and single-sex tissues (Supplementary Figure S8a, S8b), but did not include them in our study of mortality due to their relatively insignificant contribution to mortality [8] [9]. During the feature selection phase of our modeling, simply using the genes with high absolute Pearson correlation of expression levels to phenotypic age yielded better model performance than using the genes identified as differentially-expressed with age by DESeq2 [4] which is a much more commonly used pipeline in bioinformatics for differential gene expression analysis [46].

The possibility of applying our methodology to single-cell transcriptomic data to study aging at cellular resolution was explored; however, we concluded that this direction warrants an independent investigation. Single-cell RNA-seq data are substantially more complex than bulk transcriptomic profiles, as gene expression is indexed at the level of individual cells and cell types in addition to tissue and subject. As a result, such data are not a direct substitute for the bulk RNA-seq inputs used in our pipeline, and adapting them would require extensive preprocessing and modeling decisions that are not necessarily aligned with our biological objectives. One commonly used strategy is pseudo-bulking, whereby per-cell expression profiles are aggregated to generate bulk-like samples; however, this approach dramatically reduces the effective sample size, often resulting in less than 50 samples per tissue–cell-type combination even in large datasets (example provided in Supplementary Section 9). Recent studies applying both true single-cell age prediction and pseudo-bulk aggregation have reported limited performance, with single-cell models typically achieving MAEs of approximately 8–10 years and Pearson correlations around 0.2–0.5, and pseudo-bulk models showing modest improvements due to noise reduction [63] [64]. More importantly, these studies primarily focus on cell-type–specific aging, whereas our work is centered on understanding systemic versus tissue-specific aging across the human body.

Our study uncovers numerous transcriptomic biomarkers of aging in the form of model weights assigned to features, i.e., coefficients assigned to the expression values of genes. Although the reliability of many of these identified biomarkers is supported by the body of existing literature from experimental studies, a large number of them have not been explored at all. Within the list of genes corresponding to our model coefficients, there is abundant scope for future research in the field of experimental molecular biology. Such research should aim to empirically determine the correlation of the expression of specific genes with diseases of the associated tissue and ultimately mortality. As the body of human transcriptomic data grows larger, deep learning methods may also be explored in the area of transcriptomic age modeling.

### Key Points

- We developed machine learning models to predict tissue-specific biological ages that exceed the performance of existing transcriptomic age calculators. (Table S11)
- We introduce a transcriptomic age-gap metric to measure deviations in predicted biological ages relative to other samples and analyze their associations with mortality.
- We identify tissue-specific transcriptomic markers associated with aging in each type of tissue, as well as genes that universally regulate aging across multiple tissues.

## Data Availability

The data used in this study are publicly available from multiple large-scale transcriptomic resources.

Bulk tissue gene expression data (TPM) and associated metadata were obtained from the Genotype-Tissue Expression (GTEx) Project v10, available at https://www.gtexportal.org/home/downloads/adult-gtex/bulk_tissue_expression and https://www.gtexportal.org/home/downloads/adult-gtex/metadata. Access to GTEx data is subject to the terms and conditions of the GTEx Consortium.

External validation was performed using transcriptomic data from the Human Protein Atlas (HPA) Project (version 24.1), which provides RNA-seq–based transcript-level expression (TPM) across multiple human tissues. The HPA data are publicly accessible at https://www.proteinatlas.org/about/download.

Additional cross-cohort evaluation was conducted using normal lung tissue samples from The Cancer Genome Atlas Lung Adenocarcinoma (TCGA-LUAD) dataset. TCGA-LUAD expression data were accessed via the UCSC Xena platform at https://xenabrowser.net/datapages/. Only solid normal tissue samples were used in this study, consistent with model training on non-diseased tissues.

## Biographical Notes

**Md. Abul Hassan Samee** is an Associate Professor at Baylor College of Medicine. He earned his PhD in Computer Science from the University of Illinois Urbana-Champaign and completed postdoctoral training at Gladstone Institutes. His group develops machine learning methods for single-cell and spatial omics to study aging, regeneration, and disease.

**M. Sohel Rahman** is a Professor in the Department of Computer Science and Engineering at Bangladesh University of Engineering and Technology. He received his PhD from King’s College London. His research covers algorithms, bioinformatics, and machine learning.

## Supplementary Materials

ORANGE: A Machine Learning Approach for Modeling Tissue-Specific Aging from Transcriptomic Data

## 1 Dataset

### 1.1 The Hardy Scale

The Hardy Scale is a categorical measure used to assess the overall health status of individuals at the time of death, in the Genotype-Tissue Expression Project (GTEx). It provides a standardized framework for classifying subjects based on the presence and severity of disease conditions. The scale is typically assigned based on clinical and pathological evaluations and is often used in research involving aging, mortality risk, and disease progression. The scale consists of the following categories:

- **Type 1 (Violent and fast death of unnatural causes):** Deaths due to accident, blunt force trauma, or suicide, terminal phase estimated at less than 10 minutes.
- **Type 2 (Fast death of natural causes):** Sudden unexpected deaths of people who had been reasonably healthy, after a terminal phase estimated at less than 1 hour (with sudden death from a myocardial infarction as a model cause of death for this category).
- **Type 3 (Intermediate death):** Death after a terminal phase of 1 to 24 hours (not classifiable as 2 or 4); patients who were ill but death was unexpected.
- **Type 4 (Slow death):** Death after a long illness, with a terminal phase longer than 1 day (commonly cancer or chronic pulmonary disease); deaths that are not unexpected.
- **Type 0 (Ventilator case):** All cases on a ventilator immediately before death.
- In studies on the GTEx dataset, including this one, the Hardy Scale ratings of subjects are typically defined by the variable named *dthhrdy*, as done in the GTEx study.

### 1.2 Age range distribution in GTEx phenotypes

**Figure S1:**
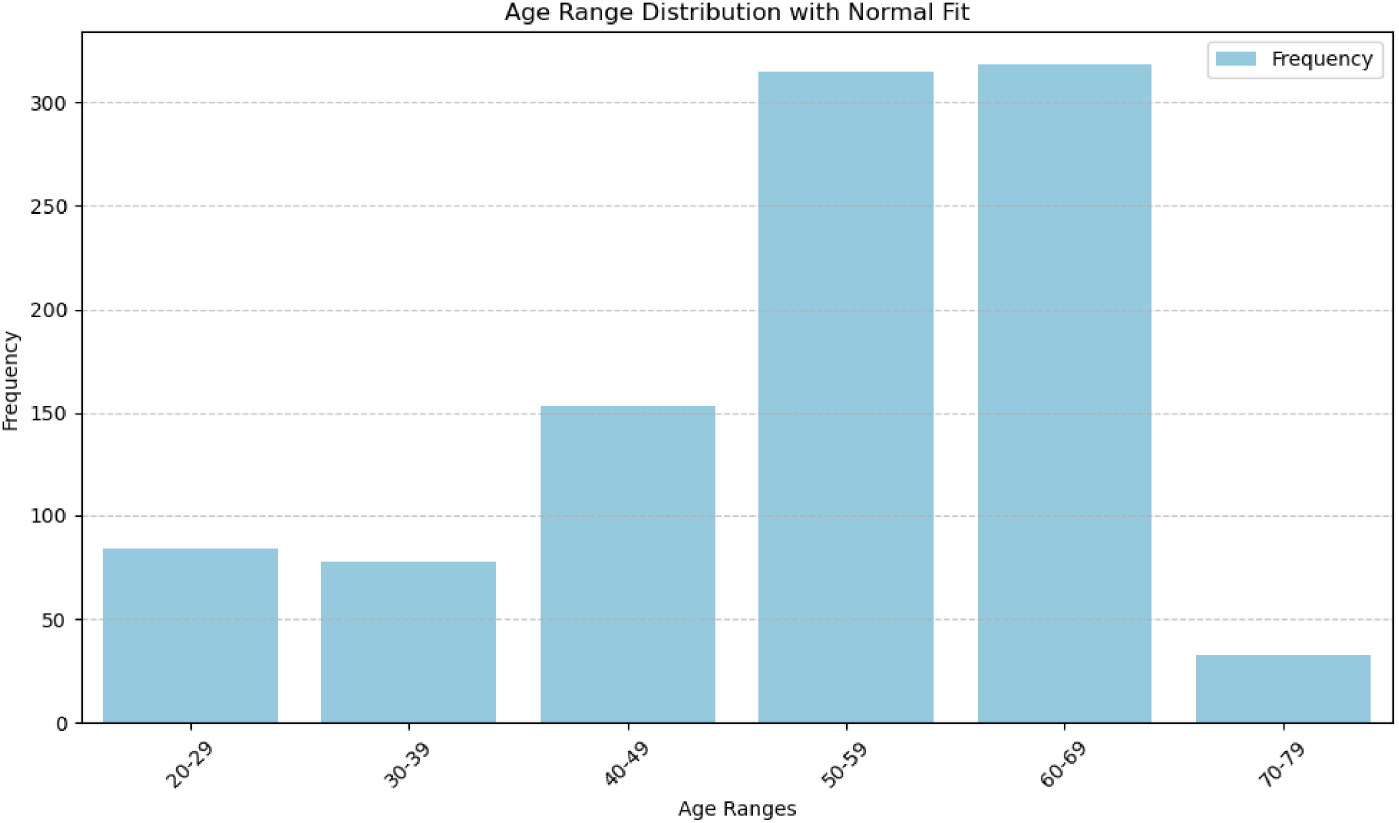
Age distribution of GTEx subjects.

## 2 Linear Regression Models

### 2.1 Least Absolute Shrinkage and Selection Operator (LASSO) Regression

LASSO regression is a linear model that adds an L1 penalty to the loss function, encouraging sparsity in feature selection. This helps in reducing overfitting by shrinking some coefficients to zero. The LASSO regression minimizes the following objective:

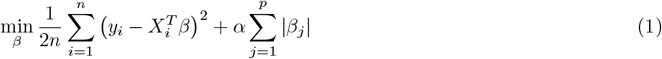

where *X* is the feature matrix, *y* is the target vector, *β* represents the regression coefficients, *n* is the number of observations, *p* is the number of features, and *α* is the regularization parameter controlling the amount of shrinkage.

### 2.2 Elastic Net Regression

Elastic Net combines LASSO (L1 penalty) and Ridge regression (L2 penalty) to balance feature selection and coefficient shrinkage. It is useful when features are correlated.Elastic net regression minimizes the following objective:

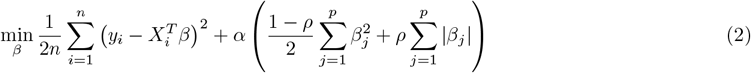

where *ρ* controls the balance between L1 and L2 regularization, and *α* remains the overall regularization strength.

### 2.3 Partial Least Squares (PLS) Regression

PLS regression [**?**] is a dimensionality reduction technique that projects predictor variables into a shared low-dimensional space to maximize covariance with the response variable. It is useful for highly collinear data or when the number of predictors exceeds the number of observations. PLS regression creates new predictor variables, called components, as linear combinations of the original predictor variables, selected to maximize the covariance with the target variable. These components are then used to predict the response variable. The PLS model assumes a latent variable approach. The observed variables *X* and *y* are described as linear combinations of unobserved latent variables plus random noise as follows:

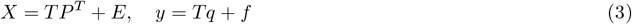

where *T* is the score matrix, *P* is the loading matrix for *X, q* is the loading vector for *y, E* and *f* are residuals, and *T* is shared between *X* and *y*, ensuring that the components represent a shared structure.

### 2.4 Support Vector Regression (SVR)

SVR aims to find a hyperplane that predicts *y* with at most *ϵ*-deviation while maximizing the margin. The objective is:

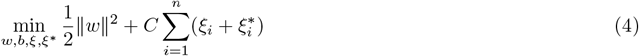

subject to

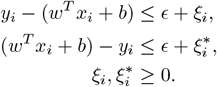

Here, *C* is the regularization parameter, w is the weight vector, b is the bias term, *ϵ* defines the margin of tolerance, and *ξ, ξ*^∗^ are slack variables.

The **linear kernel** version assumes a linear relationship between inputs and outputs.

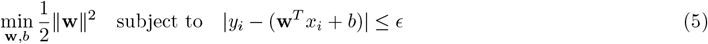

## 3 Ensemble Learning

Ensemble learning is a powerful technique in machine learning where multiple models are combined to improve predictive performance. By leveraging the strengths of different models, ensemble methods reduce variance, bias, and overfitting, leading to more robust and accurate predictions. Two popular ensemble approaches are Bootstrap Aggregating (Bagging) and Random Forest Regression, which enhance model stability and generalization.

### 3.1 Bootstrap Aggregating (Bagging)

Bagging is an ensemble method that improves accuracy and reduces variance by training multiple models on different bootstrap samples (random subsets with replacement) of the dataset. The final prediction is obtained by averaging (for regression) or majority voting (for classification).The prediction is given by:

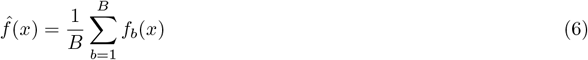

where *f*_*b*_(*x*) represents individual base models trained on different bootstrap samples, and *B* is the total number of models.

### 3.2 Random Forest Regression

Random Forest regression uses multiple decision trees trained on random subsets of data and features. It improves prediction accuracy and reduces overfitting. The final output is obtained by averaging the predictions of all trees. The prediction is given by:

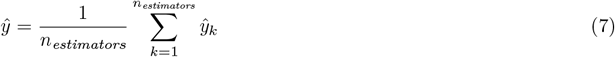

where *ŷ*_*k*_ is the prediction from the *k*-th decision tree and *n*_*estimators*_ is the number of trees in the ensemble.

## 4 Experimental setup for machine learning

To predict tissue ages, we trained LASSO, Elastic Net, SVR, PLS Regression, and Random Forest Regression predictors using the Python package scikit-learn.

Linear predictors and SVR were trained with a 0.01 tolerance for stopping criterion, and a maximum of 50,000 iterations. All predictors other than random forest regression were 20× bootstrapped. An L1-to-L2 regularization ratio of 0.5 was applied in training Elastic Net predictors.

### 4.1 Hyperparameter tuning

All predictors other than PLS regression underwent hyperparameter tuning, which was done using the GridSearchCV tool from scikit-learn. Four-fold cross validation was applied for hyperparameter tuning, using negative mean-absolute error as the scoring metric, and it significantly improved the performance of prediction algorithms, as can be seen from the following example of Elastic Net.

**Table S1:** Performance impact of cross validation.

| Tissue | Without Cross-validation |  | With Cross-validation |  |
| --- | --- | --- | --- | --- |
|  | RMSE | R<sup>2</sup> | RMSE | R<sup>2</sup> |
| Liver | 6.83 | 0.41 | 6.02 | 0.54 |
| Sigmoid Colon | 7.70 | 0.54 | 6.52 | 0.67 |
| Sun-Exposed Skin | 7.64 | 0.59 | 6.81 | 0.68 |
| Subcutaneous Adipose | 7.97 | 0.51 | 7.01 | 0.62 |
| Coronary Artery | 7.47 | 0.49 | 5.39 | 0.73 |
| Aorta | 6.05 | 0.67 | 5.32 | 0.74 |
| Brain Cortex | 7.76 | 0.42 | 6.46 | 0.59 |
| Brain Cerebellum | 8.65 | 0.31 | 7.88 | 0.43 |
| Heart Atrial Appendage | 7.95 | 0.22 | 7.23 | 0.36 |
| Lung | 8.25 | 0.42 | 6.27 | 0.66 |
| Tibial Nerve | 6.45 | 0.70 | 5.44 | 0.79 |
| Pancreas | 6.61 | 0.59 | 6.45 | 0.61 |

In order to prevent overfitting and achieve better generalization, instead of taking the best hyperparameters from grid search, a 95th percentile performance cutoff was used for selecting optimal hyperparameters.

**Figure S2:**
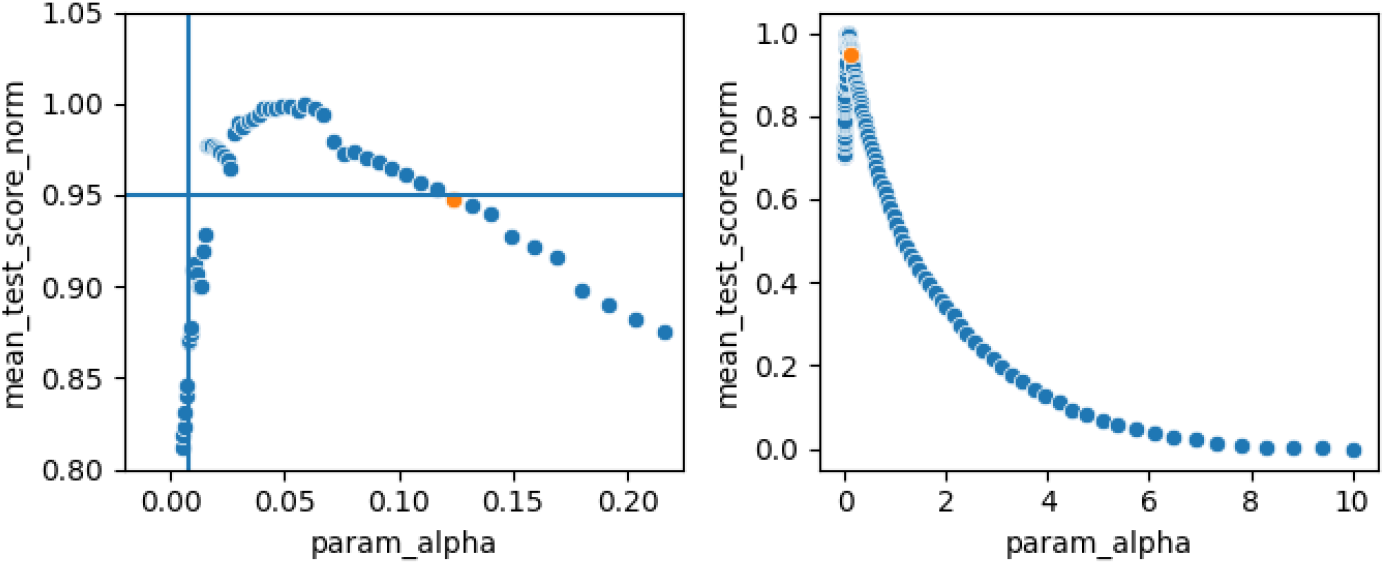
Application of performance cutoff in hyperparameter tuning.

Cross validation and hyperparameter training were skipped during the training of PLS regression predictors, because the long running time incurred would make them infeasible for the downstream tasks. PLS regression was preferred because it was observed to offer robust performance comparable to the other methods even without cross-validation (Supplementary Table S7).

However, we applied an approximation method with some hand-tuning based on repeated observations on smaller subset datasets to fix the value of the *n*_*components*_ parameter for the regressor trained on each tissue type. The approximation was *n*_*components*_ = *F* × *n*_*features*_, where F is a factor hand-tuned for each tissue type.

**Table S2:** Component decomposition factor across tissues.

| Tissue | Factor |
| --- | --- |
| Liver | 0.02 |
| Aorta | 0.10 |
| Coronary Artery | 0.02 |
| Brain Cortex | 0.01 |
| Brain Cerebellum | 0.003 |
| Adrenal Gland | 0.01 |
| Heart Atrial Appendage | 0.035 |
| Pituitary | 0.01 |
| Subcutaneous Adipose | 0.10 |
| Lung | 0.01 |
| Sun-Exposed Skin | 0.02 |
| Tibial Nerve | 0.01 |
| Sigmoid Colon | 0.035 |
| Pancreas | 0.09 |

The exact values used in tuning of hyperparameters are summarized in the following table.

**Table S2:**
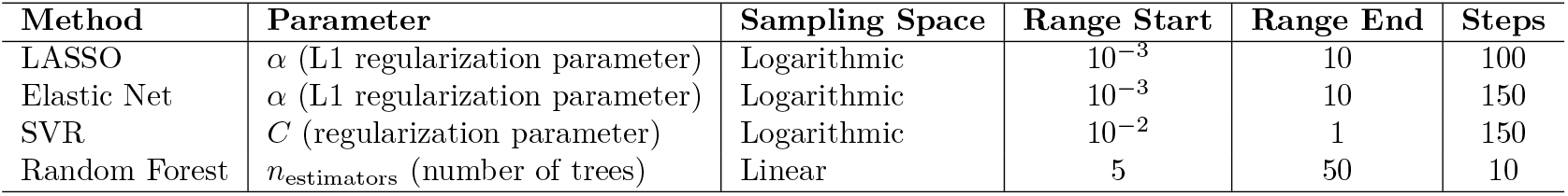
Hyperparameter search configurations.

## 5 Stratified sampling strategy

### Algorithm 1 Organ-wise data preparation and stratified splitting

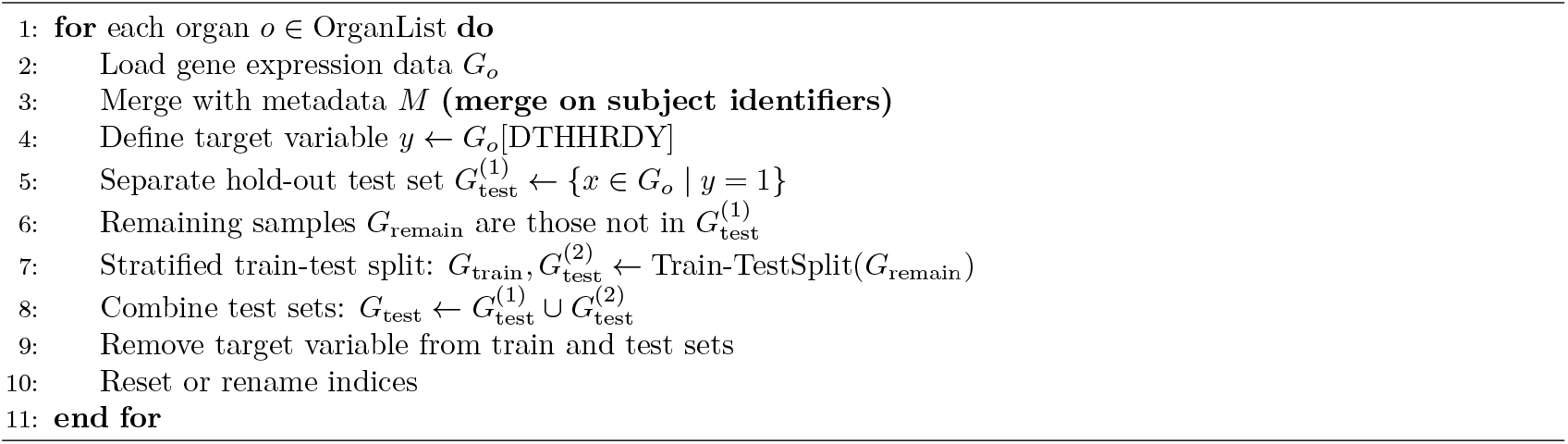

## 6 Experimental setup for cross-cohort testing

To test our models on external datasets, the collection of GTEx genes that our models were originally trained on had to be subsetted to the overlapping genes whose expression levels were available in each external cohort. Models were retrained on the new subset of columns and then tested on external cohort data.

### 6.1 The Human Protein Atlas (HPA) Project

The Human Protein Atlas (HPA) transcriptomics dataset provides a comprehensive view of gene expression across a wide range of human tissues. In version 24.1, transcript-level RNA-seq data are available for 186 distinct tissue samples, encompassing 40,539 protein-coding genes. The dataset captures normalized expression levels in transcripts per million (TPM), offering a consistent framework for comparing transcript abundance across tissues. By integrating these data, the HPA serves as a valuable resource for exploring tissue-specific transcriptional patterns. Implementation details for testing on this dataset are provided along with performance metrics in the following table.

**Table S2:** Performance of transcriptomic age prediction models across tissues.

| Tissue Type |  | Original GTEx-Trained Model |  |  | Subset-Trained Model (Val. on GTEx) |  |  | Test on HPA |  |  |
| --- | --- | --- | --- | --- | --- | --- | --- | --- | --- | --- |
| GTEx | HPA | $n_{\text{features}}$ | RMSE (yrs) | $R^2$ | $n_{\text{features}}$ | RMSE (yrs) | $R^2$ | RMSE (yrs) | $R^2$ | $n_{\text{samples}}$ |
| Subcut. Adipose | Adipose (unspecified) | 2191 | 7.01 | 0.62 | 1802 | 6.36 | 0.67 | 9.51 | 0.46 | 5 |
| Lung | Lung | 2019 | 6.27 | 0.66 | 1765 | 6.66 | 0.62 | 5.31 | 0.32 | 9 |
| Liver | Liver | 2712 | 6.02 | 0.54 | 2386 | 8.09 | 0.28 | 9.60 | -0.001 | 8 |
| Kidney Cortex | Kidney (unspecified) | 1893 | - | - | 932 | - | - | 8.32 | 0.05 | 8 |
| **Average** |  |  |  |  |  |  |  | 8.16 | 0.30 | 30 |

**Figure S3:**
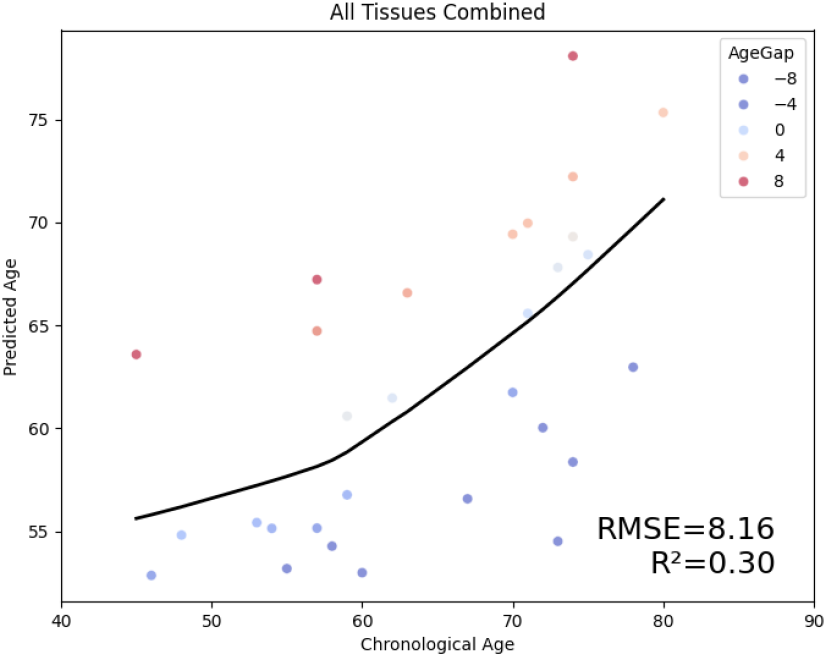
Test performance on HPA data. Model performance on 30 samples covering 4 tissues from the Human Protein Atlas project, in terms of RMSE and *R*^2^ scores.

### 6.2 The Cancer Genome Atlas — Lung Adenocarcinoma (TCGA-LUAD) Dataset

The TCGA-LUAD (The Cancer Genome Atlas – Lung Adenocarcinoma) dataset provides transcriptomic profiles for lung adenocarcinoma tumor samples and matched normal tissue. RNA-seq–derived expression levels are typically reported in transcripts per million (TPM) or raw counts, covering 60,498 protein-coding genes across several hundred patient samples. This dataset captures both tumor and adjacent normal tissue expression patterns, enabling analysis of tissue-specific gene regulation, tumor heterogeneity, and validation of predictive models developed on independent datasets. TCGA-LUAD serves as an important resource for benchmarking transcriptomic age prediction models in lung tissue and for exploring gene-level associations with disease states. For the purpose of this study we only chose the normal tissue samples since our models are trained on GTEx tissues labeled as normal. Implementation details for testing on this dataset are provided along with performance metrics in the following table.

**Table S3:**
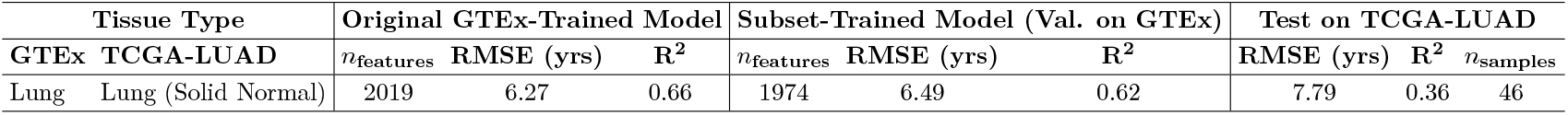
Performance of transcriptomic age prediction models on TCGA-LUAD samples.

| Tissue Type |  | Original GTEx-Trained Model |  |  | Subset-Trained Model (Val. on GTEx) |  |  | Test on TCGA-LUAD |  |  |
| --- | --- | --- | --- | --- | --- | --- | --- | --- | --- | --- |
| GTEx | TCGA-LUAD | $n_{\text{features}}$ | RMSE (yrs) | $R^2$ | $n_{\text{features}}$ | RMSE (yrs) | $R^2$ | RMSE (yrs) | $R^2$ | $n_{\text{samples}}$ |
| Lung | Lung (Solid Normal) | 2019 | 6.27 | 0.66 | 1974 | 6.49 | 0.62 | 7.79 | 0.36 | 46 |

**Figure S4:**
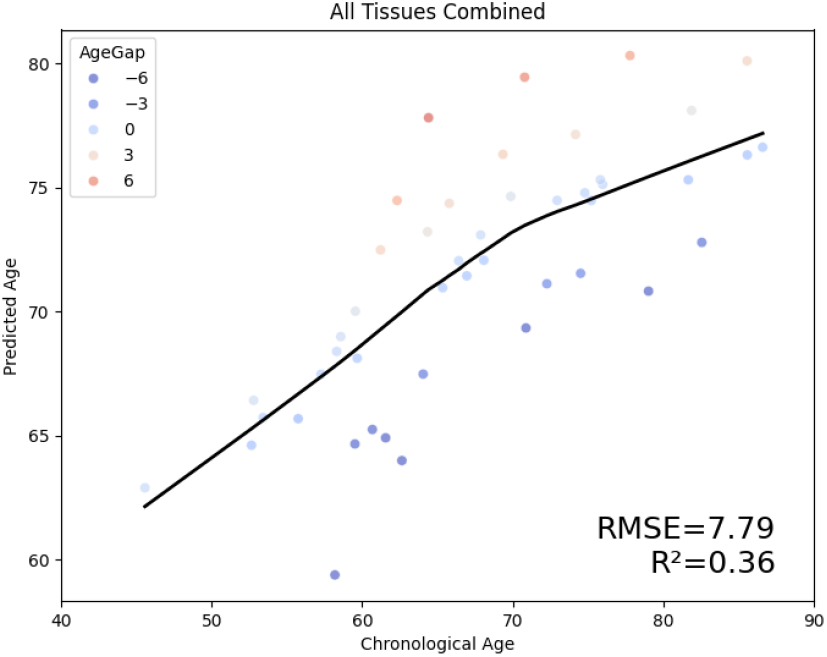
Test performance on TCGA-LUAD data. Model performance on 46 samples of normal tissue from the Lung Adenocarcinoma dataset the Cancer Genome Atlas, in terms of RMSE and *R*^2^ scores.

## 7 Clustering of gene expression (TPM) data of the samples from 12 types of tissues

**Figure S5:**
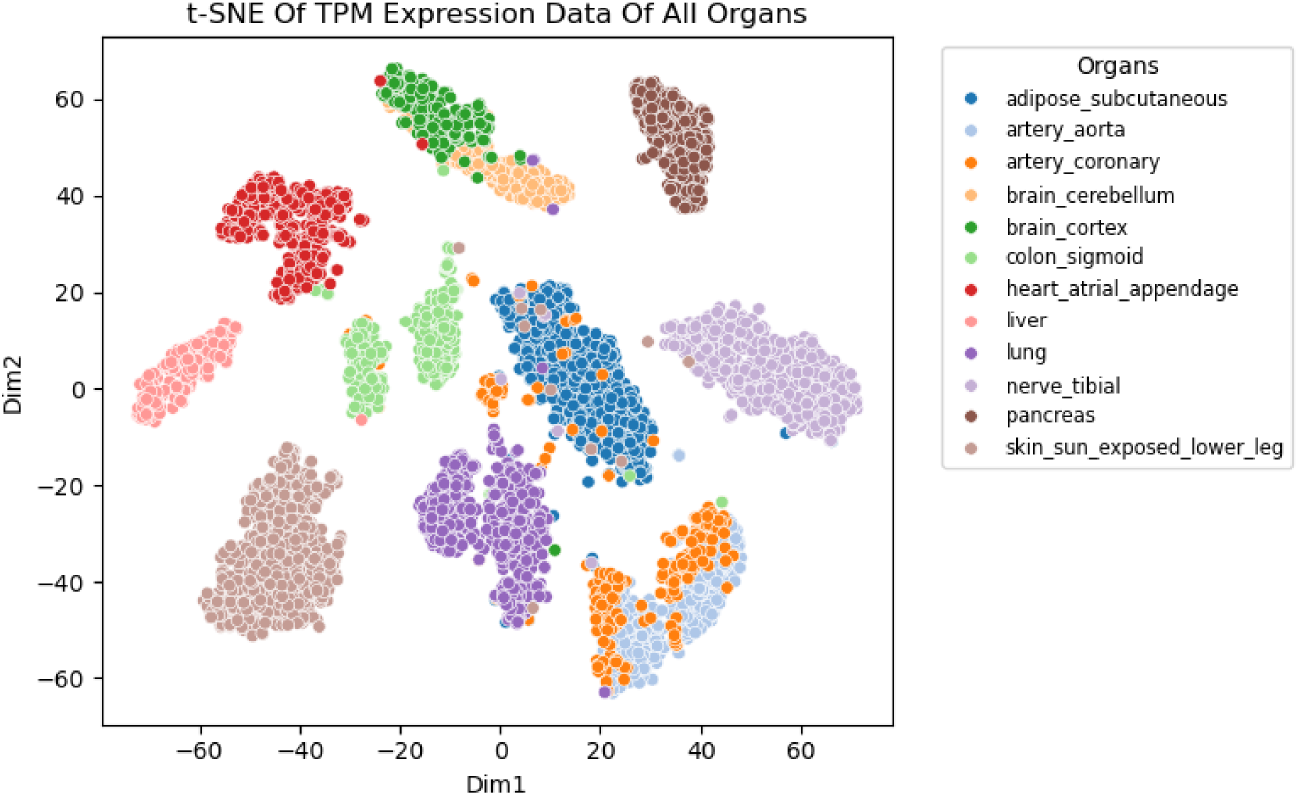
T-SNE plot of the tissue-specific gene expression (TPM) values. The two axes, Dim1 and Dim2, are low-dimensional embeddings learned by the t-SNE algorithm to best preserve local structure and relationships between high-dimensional gene expression profiles. Dots represent the RNA-Seq samples of the 12 tissue types from the Adult GTEx that we have used in our study. It is observed that the samples of the same type of tissue strongly tend to cluster together with themselves while remaining noticeably separate from the samples of other tissue types. It is also evident that tissues from the same or related organs, such as the coronary artery and aorta, or, the brain’s cerebellum and cortex, cluster adjacent to each other.

## 8 Clustering of tissue-specific gene expression (TPM) data

**Figure S6:**
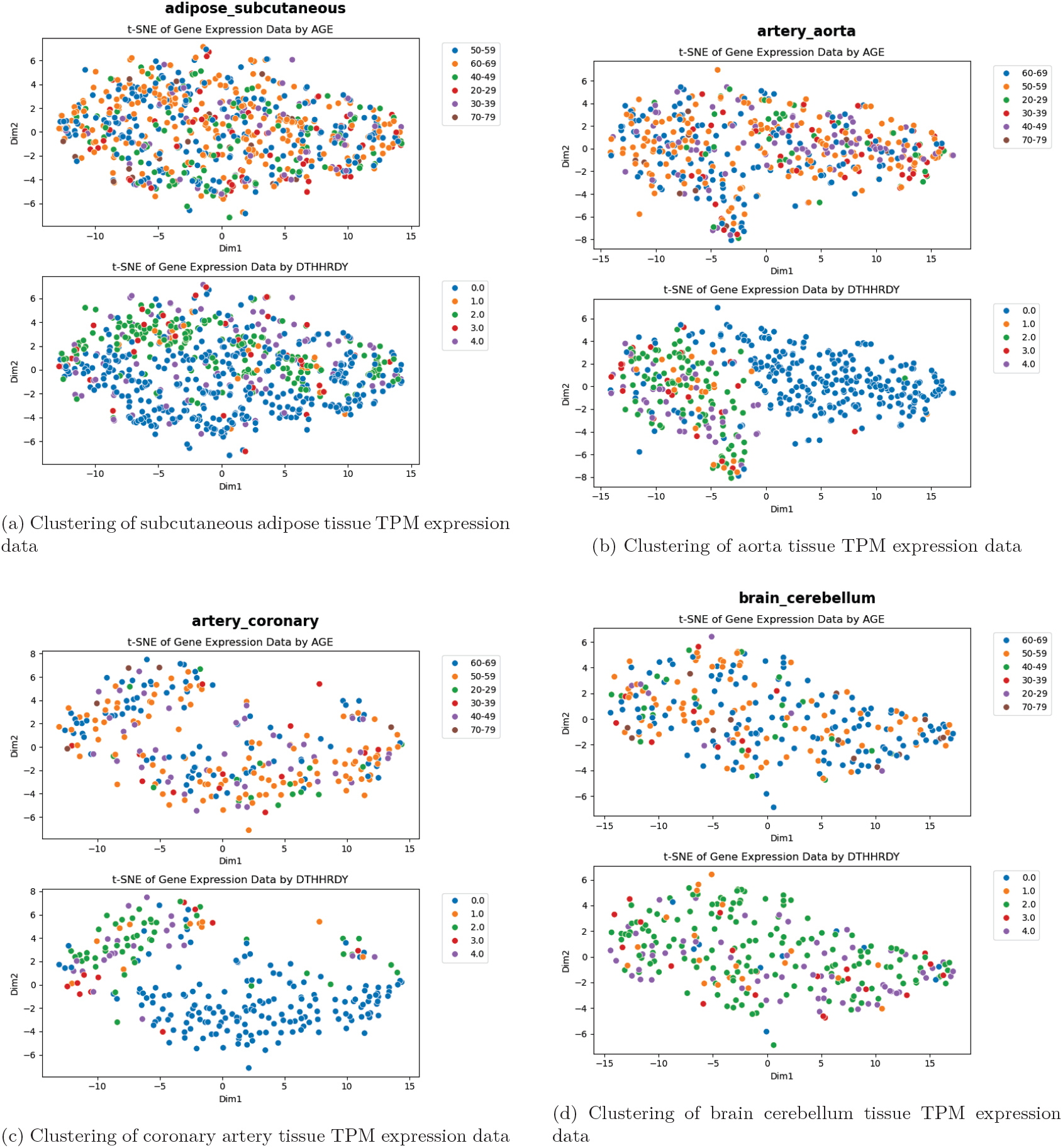
Part 1: t-SNE plots of gene expression (TPM) for individual tissues (a–d). In most tissues, it is observed that the samples from subjects with a Hardy Scale rating of 0 (death on a ventilator) tend to cluster together with themselves and separate from the samples from subjects with other death types. No significant grouped clustering by age range can be observed.

**Figure S6:**
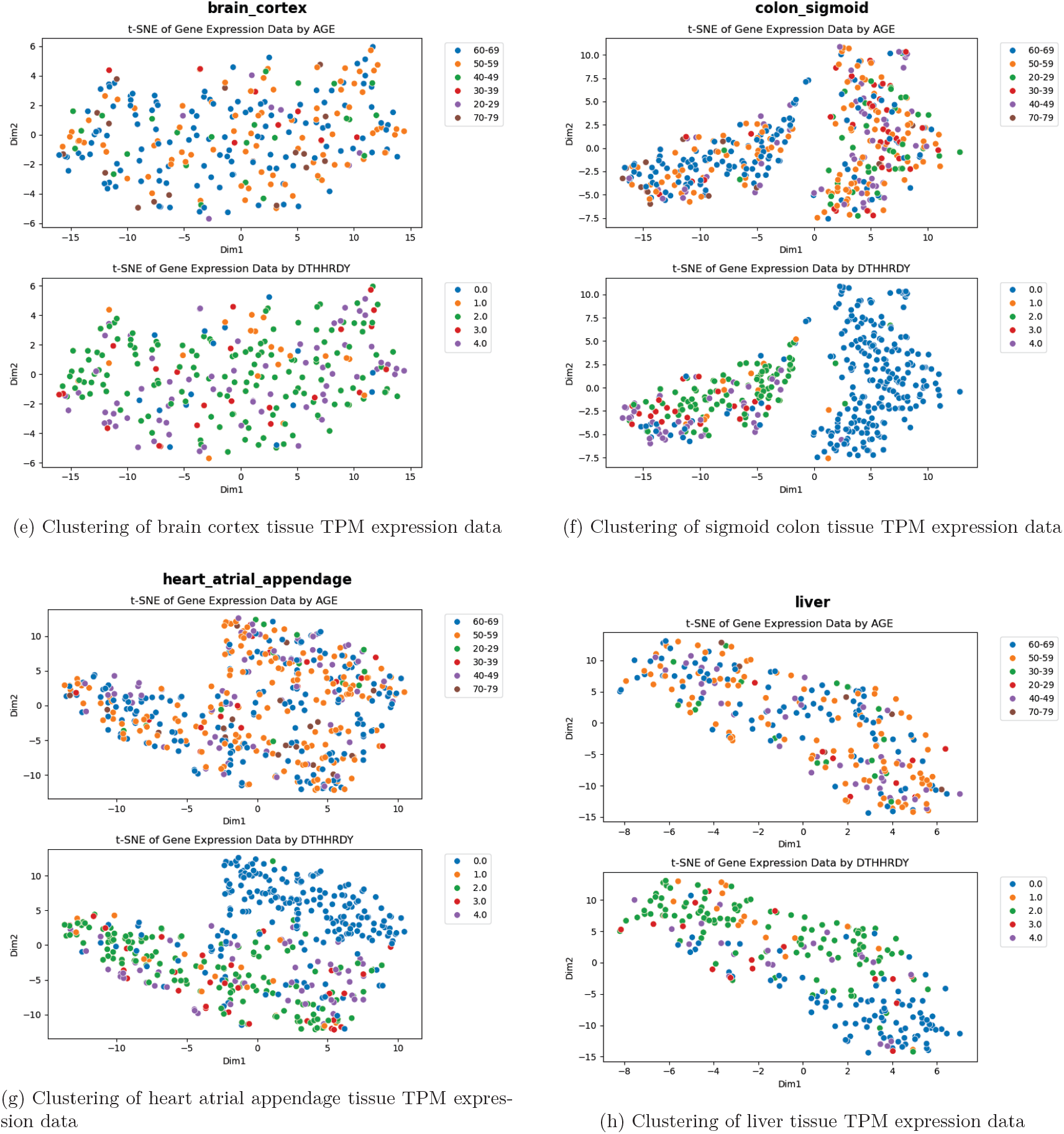
Part 2: t-SNE plots of gene expression (TPM) for individual tissues (e–h). In most tissues, it is observed that the samples from subjects with a Hardy Scale rating of 0 (death on a ventilator) tend to cluster together with themselves and separate from the samples from subjects with other death types. No significant grouped clustering by age range can be observed.

**Figure S6:**
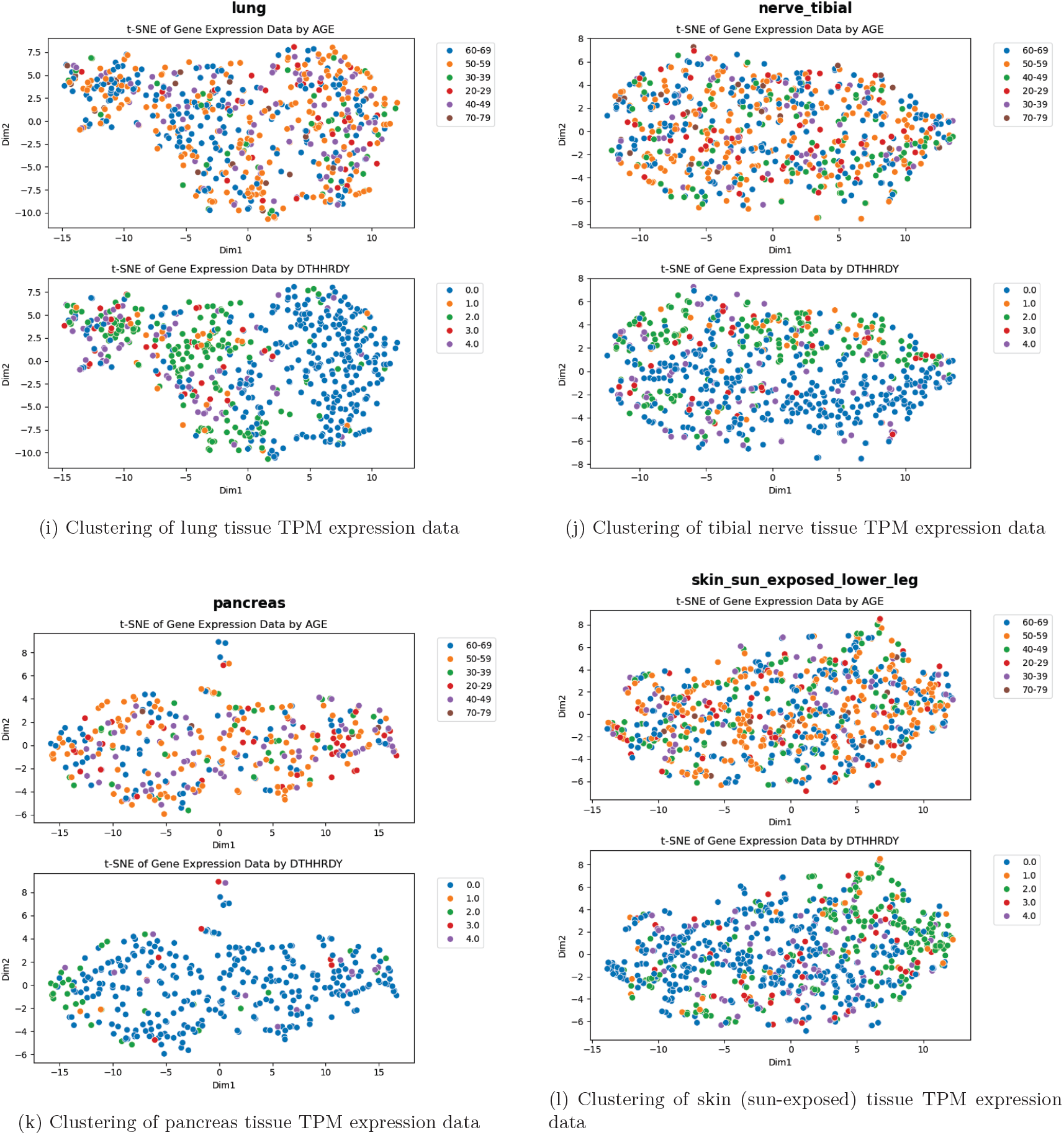
Part 3: t-SNE plots of gene expression (TPM) for individual tissues (i–l). In most tissues, it is observed that the samples from subjects with a Hardy Scale rating of 0 (death on a ventilator) tend to cluster together with themselves and separate from the samples from subjects with other death types. No significant grouped clustering by age range can be observed.

## 9 Single-Cell Data Availability for Tissue-Specific Age Modeling

The Human Cell Atlas Tissue Atlas (HCATA) is one of the largest publicly available age-annotated single-cell transcriptomic resources and among the few datasets that provide cell-type–resolved gene expression profiles across multiple human tissues. HCATA integrates single-cell RNA sequencing data from a diverse set of organs and annotates cells by tissue of origin and cell type, making it a valuable resource for studying cellular heterogeneity across tissues. However, despite its scale relative to other single-cell resources, HCATA remains limited in terms of the number of samples available per tissue–cell-type combination, particularly when contrasted with large bulk transcriptomic datasets such as GTEx.

Table S6 summarizes the availability of single-cell samples from HCATA for tissue types common to both HCATA and GTEx and included in our study, alongside the corresponding number of bulk RNA-seq samples available in GTEx.

**Table S6:**
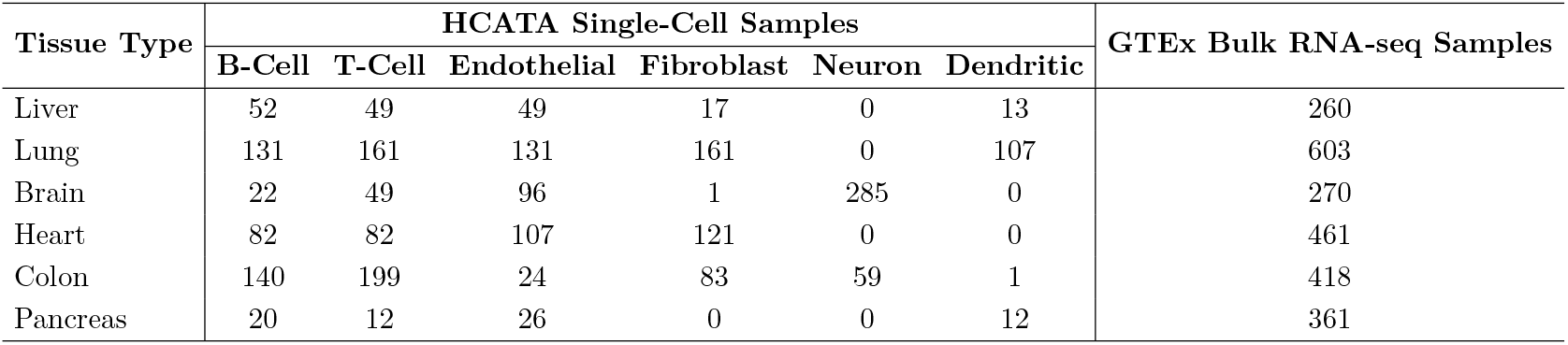
Comparison of Single-Cell and Bulk Transcriptomic Data Availability Across Tissues.

| Tissue Type | HCATA Single-Cell Samples |  |  |  |  |  | GTEx Bulk RNA-seq Samples |
| --- | --- | --- | --- | --- | --- | --- | --- |
|  | B-Cell | T-Cell | Endothelial | Fibroblast | Neuron | Dendritic |  |
| Liver | 52 | 49 | 49 | 17 | 0 | 13 | 260 |
| Lung | 131 | 161 | 131 | 161 | 0 | 107 | 603 |
| Brain | 22 | 49 | 96 | 1 | 285 | 0 | 270 |
| Heart | 82 | 82 | 107 | 121 | 0 | 0 | 461 |
| Colon | 140 | 199 | 24 | 83 | 59 | 1 | 418 |
| Pancreas | 20 | 12 | 26 | 0 | 0 | 12 | 361 |

As shown in Table S6, the volume of single-cell data available per tissue and per cell type is significantly smaller than the corresponding bulk RNA-seq sample sizes in GTEx. This scarcity severely limits the feasibility of meaningful linear modeling at cellular resolution for tissue-specific age prediction. Furthermore, these counts do not account for the requirement of splitting data into non-overlapping subject-level training and validation sets to avoid data leakage in true single-cell modeling. Enforcing such splits further reduces the effective sample size, often to impractically small levels. Pseudo-bulking approaches exacerbate this issue by merging cells from the same subject, resulting in even fewer data points. Consequently, despite HCATA being among the largest and most comprehensive tissue-resolved single-cell transcriptomic datasets currently available, it does not provide sufficient data to support robust age modeling in a manner that is both statistically reliable and aligned with the biological objectives of our study.

## 10 Optimal point sampling strategy

**Figure S7:**
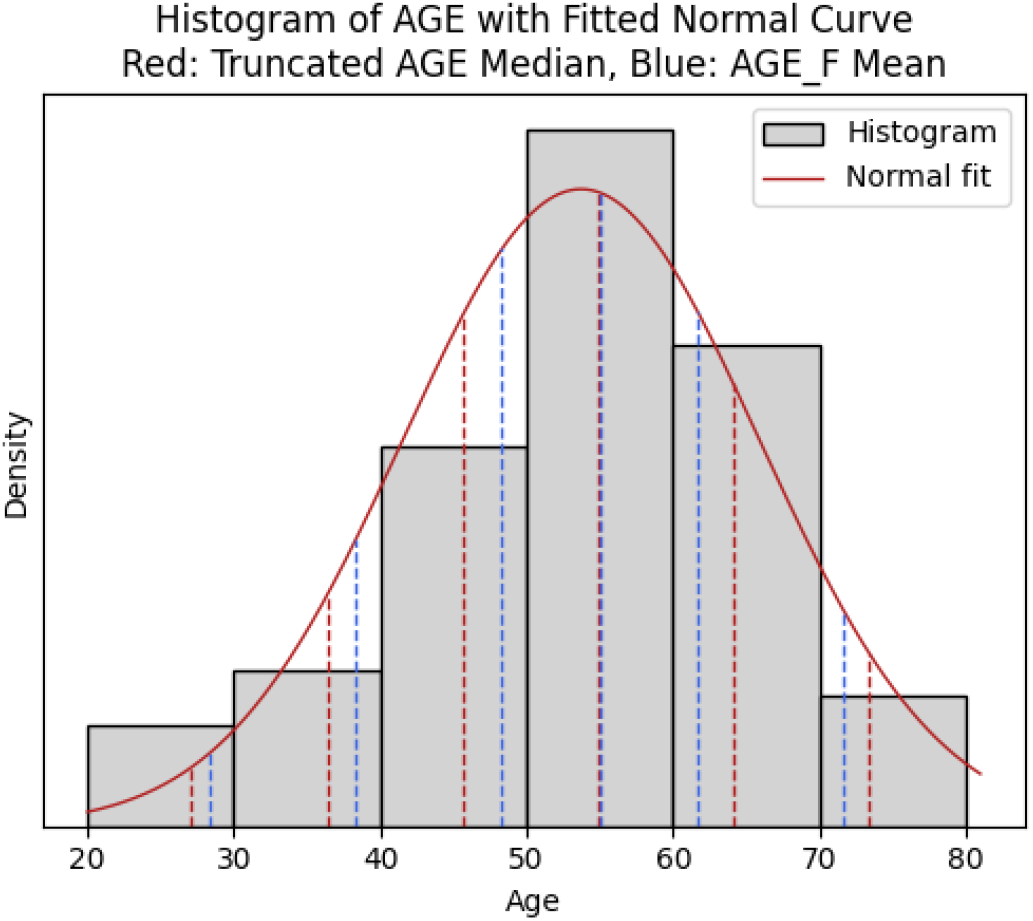
Motivation for optimal age sampling strategy. We assume that the ages of subjects are expected to follow a normal distribution. When we fit a normal curve to the distribution, it is observed that the expected mean of the portion of the normal curve within each age range (indicated by dashed red lines) can be deviated from the mid-point towards the center of the curve, i.e. the mean of the entire distribution. The combination “222100” is a recurring top result produced by our sampling method, and plotting the age points corresponding to that combination (indicated by blue dashed lines), we notice that they are also deviated towards the sample mean, suggesting that the points we call ‘optimal’ represent a more accurate picture of the age distribution than what is achieved by directly taking mid-points.

**Table S7:** Performance impact of optimal fixed-point sampling method. The fixed-point configuration produced by our sampling method, was found to consistently achieve lower RMSE scores and higher *R*^2^ scores across 7 tissue types. Through an exhaustive search we found that it also consistently ranked high among all 3^6^ configurations based on RMSE.

| Tissue Type | Configuration “111111”<br>(All midpoints) | | Configuration “222100”<br>(Chosen as optimal) | | Rank of configuration<br>“222100” out of $3^6$<br>RMSE Rank |
| --- | --- | --- | --- | --- | --- |
| | RMSE | $R^2$ | RMSE | $R^2$ | |
| Liver | 6.15 | 0.5203 | 6.08 | 0.5297 | 2 |
| Aorta | 6.84 | 0.7202 | 5.47 | 0.7403 | 4 |
| Coronary Artery | 5.96 | 0.7104 | 5.24 | 0.7299 | 1 |
| Heart Atr. App. | 7.81 | 0.3901 | 5.90 | 0.3957 | 3 |
| Lung | 7.93 | 0.6398 | 6.46 | 0.6602 | 1 |
| Sun-Exposed Skin | 8.42 | 0.6699 | 7.02 | 0.6797 | 7 |
| Pancreas | 7.64 | 0.5897 | 6.26 | 0.6102 | 2 |

## 11 Conditional probability analysis of maximum age-gap of all subjects

**Table S7:** Conditional probabilities of DTHHRDY given different age-gap ranges.

| DTHHRDY | Agegap $\geq 1.5353$ | Agegap $\leq 0.8033$ | $1.0208 < \text{Agegap} < 1.3148$ | Overall ($p$) |
| --- | --- | --- | --- | --- |
| 0 | 0.3377 | 0.6980 | 0.3864 | 0.5066 |
| 1 | 0.0265 | 0.0336 | 0.0568 | 0.0377 |
| 2 | 0.3974 | 0.1879 | 0.3636 | 0.2994 |
| 3 | 0.0662 | 0.0201 | 0.0682 | 0.0508 |
| 4 | 0.1722 | 0.0604 | 0.1250 | 0.1055 |

The following observations were made on the basis of the calculated conditional probabilities:

- ***dthhrdy*=0**:
  - Extreme negative agers are 2.067 times as likely to have died with *dthhrdy*=0 compared to extreme positive agers.
  - Average agers are 1.144 times as likely to have died with *dthhrdy*=0 compared to extreme positive agers.
  - Extreme negative agers are 1.807 times as likely to have died with *dthhrdy*=0 compared to average agers.
- ***dthhrdy*=1**:
  - Extreme negative agers are 1.267 times as likely to have died with *dthhrdy*=1 compared to extreme positive agers.
  - Average agers are 2.145 times as likely to have died with *dthhrdy*=1 compared to extreme positive agers.
  - Average agers are 1.693 times as likely to have died with *dthhrdy*=1 compared to extreme negative agers.
- ***dthhrdy*=2**:
  - Extreme positive agers are 2.114 times as likely to have died with *dthhrdy*=2 compared to extreme negative agers.
  - Extreme positive agers are 1.093 times as likely to have died with *dthhrdy*=2 compared to average agers.
  - Average agers are 1.935 times as likely to have died with *dthhrdy*=2 compared to extreme negative agers.
- ***dthhrdy*=3 or 4**:
  - Extreme positive agers are 2.96 times as likely to have died with *dthhrdy*=3 or 4 compared to extreme negative agers.
  - Extreme positive agers are 1.234 times as likely to have died with *dthhrdy*=3 or 4 compared to average agers.
  - Average agers are 2.399 times as likely to have died with *dthhrdy*=3 or 4 compared to extreme negative agers.

## 12 Performance comparison of pipeline variants

**Table S7:** Performance comparison of pipeline variants.

| Tissue Type | Metric | Elastic Net |  | PLS |  | SVR |  | Rand. Forest |  | Lasso |
| --- | --- | --- | --- | --- | --- | --- | --- | --- | --- | --- |
|  |  | Corr. | DEG | Corr. | DEG | Corr. | DEG | Corr. | DEG | Enriched |
| Liver | RMSE | 6.02 | 6.78 | 6.41 | 6.81 | 6.41 | 6.86 | 8.14 | 8.32 | 8.75 |
| | $R^2$ | 0.54 | 0.42 | 0.48 | 0.42 | 0.48 | 0.41 | 0.17 | 0.13 | 0.04 |
| Aorta | RMSE | 5.32 | 5.62 | 5.48 | 6.01 | 5.51 | 6.02 | 6.43 | 6.00 | 8.91 |
| | $R^2$ | 0.74 | 0.71 | 0.73 | 0.67 | 0.72 | 0.67 | 0.62 | 0.67 | 0.28 |
| Coronary Artery | RMSE | 5.39 | 6.81 | 5.40 | 7.06 | 5.47 | 7.19 | 8.23 | 8.65 | 10.27 |
| | $R^2$ | 0.73 | 0.57 | 0.73 | 0.54 | 0.73 | 0.53 | 0.38 | 0.31 | 0.03 |
| Brain Cortex | RMSE | 6.46 | 6.23 | 6.43 | 6.06 | 6.47 | 6.11 | 8.42 | 8.10 | 7.93 |
| | $R^2$ | 0.59 | 0.62 | 0.60 | 0.64 | 0.59 | 0.64 | 0.31 | 0.36 | 0.39 |
| Brain Cerebellum | RMSE | 7.88 | 7.08 | 8.36 | 7.21 | 8.39 | 6.94 | 8.44 | 8.02 | 9.67 |
| | $R^2$ | 0.43 | 0.54 | 0.36 | 0.52 | 0.35 | 0.56 | 0.34 | 0.41 | 0.14 |
| Heart Atr. App. | RMSE | 7.23 | 6.44 | 6.99 | 6.48 | 6.63 | 6.49 | 8.33 | 8.43 | 8.63 |
| | $R^2$ | 0.36 | 0.49 | 0.40 | 0.48 | 0.46 | 0.48 | 0.15 | 0.13 | 0.09 |
| Subcut. Adipose | RMSE | 7.01 | 7.57 | 7.38 | 7.89 | 7.42 | 7.91 | 8.66 | 9.25 | 10.31 |
| | $R^2$ | 0.62 | 0.56 | 0.58 | 0.52 | 0.58 | 0.52 | 0.42 | 0.34 | 0.18 |
| Lung | RMSE | 6.27 | 7.46 | 6.41 | 7.49 | 6.41 | 7.80 | 9.03 | 8.99 | 9.55 |
| | $R^2$ | 0.66 | 0.52 | 0.65 | 0.52 | 0.65 | 0.48 | 0.30 | 0.31 | 0.22 |
| Sun-Exposed Skin | RMSE | 6.81 | 6.12 | 6.76 | 6.69 | 7.45 | 6.70 | 8.73 | 9.02 | 9.57 |
| | $R^2$ | 0.68 | 0.74 | 0.68 | 0.69 | 0.61 | 0.69 | 0.47 | 0.43 | 0.36 |
| Tibial Nerve | RMSE | 5.44 | 6.51 | 5.69 | 6.50 | 5.69 | 6.51 | 7.51 | 7.95 | 9.26 |
| | $R^2$ | 0.79 | 0.70 | 0.77 | 0.70 | 0.77 | 0.70 | 0.60 | 0.55 | 0.39 |
| Sigmoid Colon | RMSE | 6.52 | 7.45 | 7.68 | 8.00 | 7.70 | 8.00 | 8.44 | 8.48 | 10.90 |
| | $R^2$ | 0.67 | 0.57 | 0.54 | 0.50 | 0.54 | 0.50 | 0.44 | 0.44 | 0.07 |
| Pancreas | RMSE | 6.45 | 5.88 | 7.20 | 6.39 | 7.15 | 6.38 | 6.59 | 8.35 | 10.84 |
| | $R^2$ | 0.61 | 0.67 | 0.51 | 0.61 | 0.52 | 0.62 | 0.59 | 0.34 | -0.11 |

### 12.1 Training time comparison of learning algorithms

**Table S7:** Training time (in seconds) comparison of learning algorithms.

| Tissue Type | Lasso | Elastic Net | SVR | PLS | Random Forest |
| --- | --- | --- | --- | --- | --- |
| Liver | 30 | 48 | 41 | 4 | 2 |
| Sigmoid Colon | 166 | 637 | 153 | 12 | 7 |
| Sun-Exposed Skin | 107 | 56 | 127 | 0 | 3 |
| Subcutaneous Adipose | 34 | 1526 | 237 | 27 | 10 |
| Coronary Artery | 8 | 11 | 16 | 0 | 1 |
| Aorta | 9 | 199 | 250 | 41 | 9 |
| Brain Cortex | 30 | 50 | 47 | 1 | 3 |
| Brain Cerebellum | 24 | 300 | 122 | 3 | 8 |
| Heart Atrial Appendage | 70 | 57 | 26 | 0 | 0 |
| Lung | 126 | 3151 | 145 | 2 | 6 |
| Tibial Nerve | 137 | 91 | 415 | 5 | 12 |
| Pancreas | 28 | 2 | 16 | 0 | 0 |
| **Total** | **769** | **6128** | **1595** | **95** | **61** |

\* All predictors except those using random forest regression were 20× bootstrapped.

### 12.2 Performance comparison with other studies

**Table S7:** Performance comparison with other studies.

| Study | Tissue/Organ | Model | MAE | RMSE | $R^2$ |
| --- | --- | --- | --- | --- | --- |
| Ribeiro et al. [?] | Lung | EN* | 6.29 | 8.33 | 0.59 |
|  |  | GBT* | 7.07 | 8.64 | 0.56 |
| Ours* |  | EN* | **3.71** | **5.83** | **0.63** |
| Ours* |  | PLS | 4.85 | 6.11 | 0.59 |
| Ren et al. [?] | Liver | RAC* | 9.49 | 11.71 | -0.2 |
| Ours* |  | PLS | **5.39** | **6.94** | **0.31** |
| Ren et al. [?] | Tibial Nerve | RAC* | 10.61 | 12.59 | 0.08 |
| Ours* |  | PLS | **4.71** | **5.94** | **0.70** |

\* “Ours” refers to methods proposed in this study.
\* EN, GBT, and RAC refer to the Elastic net, Gradient Boosted Trees, and RNAAgeCalc [?] models respectively.

### 12.3 Prediction models for reproductive and single-sex tissues

**Figure S8:**
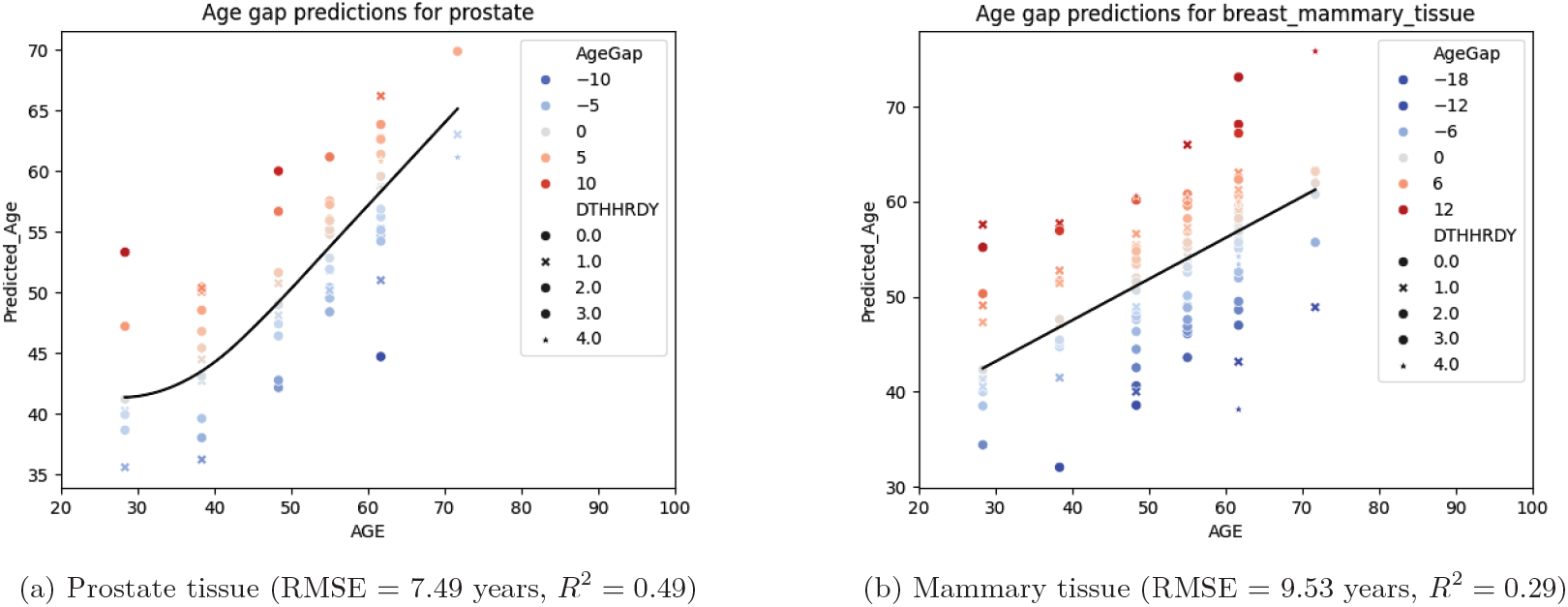
Age prediction performance in single-sex tissues.

